# An EEG study on the effect of being overweight on anticipatory and consummatory reward in response to pleasant taste stimuli

**DOI:** 10.1101/2021.09.15.460451

**Authors:** Stephanie Baines, Imca S. Hensels, Deborah Talmi

**Author notes:** Corresponding author: Stephanie Baines. Author contributions: IH, DT, and SB designed the study; IH collected the data; IH analysed the data, SB and IH wrote the paper.

## Abstract

Two-thirds of adults in the United Kingdom currently suffer from overweight or obesity, making it one of the biggest contributors to health problems. Within the framework of the incentive sensitisation theory, it has been hypothesised that overweight people experience heightened reward anticipation when encountering cues that signal food, such as pictures and smells of food, but that they experience less reward from consuming food compared to normal-weight people. There is, however, little evidence for this prediction. Few studies test both anticipation and consumption in the same study, and even fewer with electroencephalography (EEG). This study sought to address this gap in the literature by measuring scalp activity when overweight and normal-weight people encountered cues signalling the imminent arrival of pleasant and neutral taste stimuli, and when they received these stimuli. The behavioural data showed that there was a smaller difference in valence ratings between the pleasant and neutral taste in the overweight than normal-weight group, in accordance with our hypothesis. However, contrary to our hypothesis, the groups did not differ in their electrophysiological response to taste stimuli. Instead, there was a reduction in N1 amplitude to both taste and picture cues in overweight relative to normal-weight participants. This suggests that reduced attention to cues may be a crucial factor in risk of overweight.

## 1. Introduction

In the United Kingdom, 67% of adults are currently overweight or obese (World Health Organization, 2016). This poses a financial strain on health services (Davies et al., 2016) as well as a burden on people’s mental (Carr & Friedman, 2005; Falkner et al., 2001; Jackson et al., 2015) and physical health (Mokdad et al., 2003; Must et al., 1999). Prior research investigating neural differences in overweight and normal-weight individuals suggests there may be changes in reward processes relating to food. These differences can be observed both in response to food itself, and in the anticipated reward when expecting food delivery. One prominent theory, the Dynamic Vulnerability Theory and its revised form (R-DVM) (Stice et al., 2011, Burger & Stice, 2011; Stice & Yokum, 2016), suggests that obesity is characterised by an expcreased reward anticipation and a reduced experienced reward at the time of consumption. We sought to test this theory by charting the time-course of processing in overweight and normal-weight participants in expectation and upon delivery of pleasant and neutral taste stimuli.

To understand why obesity might develop, it is important to gain an understanding of the neural changes that occur in overweight and normal-weight individuals. Such differences are evident response to anticipation of food. Food cues are stimuli that signal the availability of food, such as images or smells of food (Blechert et al., 2014). Through learned association with a rewarding outcome, such as satiation of hunger or the pleasure of a tasty meal, these cues themselves acquire reward value, termed ‘incentive salience’. Cues with high incentive salience capture attention and trigger craving and consumption of food (Berridge, 2009). Some individuals experience hypersensitivity to cues, triggering heightened anticipated reward, craving, preoccupation and (food) seeking behaviour decoupled from physiological need (Berridge, 2009; Everitt & Robbins, 2005; Robinson & Berridge, 1993; Volkow & Wise, 2005). The Incentive Salience model proposes that weight gain and eventual obesity are a result of increased inscentive salience of cues, which elicit excessive intake in the absence of hunger (Appelhans, 2009; Hendrikse et al., 2015). These predicted differences in anticipation are supported by fMRI evidence. When anticipating delivery of a palatable taste stimulus, obese girls exhibit more activation compared to normal-weight girls in the insula and the frontal operculum, areas related to the processing of gustatory stimuli (Stice et al., 2008). Moreover, presentation of cues in the form of pictures of high-calorie, palatable foods is associated with increased activation in obese participants in the insula, as well as orbitofrontal cortex (OFC), striatum, and amygdala, areas associated with the processing of reward and emotionally charged stimuli (Stice & Burger, 2019; García-García, Jurado, et al., 2013). A recent meta-analysis found that this difference between obese and normal-weight individuals was greater when participants were satiated (Devoto et al., 2018). There is also evidence to suggest that heightened sensitivity to food cues predisposes obese people to overconsume after exposure to food cues (Lawrence et al., 2012; Stice & Burger, 2019). Taken together, there is consistent evidence that obese people show enhanced response to food cues compared to normal-weight people, and that this may lead to overeating.

Neural responses to receipt of food itself have also been compared in overweight and normal-weight individuals. There is evidence for differences in the neural responses in obese participants, but the nature of these differences is inconsistent and even contradictory. Some studies observe increased activation in response to delivery of palatable food, such as milkshakes, particularly in regions of the reward network, such as the ventral tegmental area (VTA), ventral striatum, orbitofrontal cortex (OFC) and insula (Stice et al., 2008b; Davis et al., 2004,. Others observe the opposite, a decrease in reward response to food delivery (Babbs et al., 2013; Green et al., 2011; Stice et al., 2008; Stice & Burger, 2019; Volkow et al., 2008), consistent with the downregulation of dopamine D2 receptors with increasing BMI (Wang et al., 2002). The latter pattern of results suggest overweight people might have less capacity for dopamine uptake in the striatum following exposure to a rewarding experience, which could result in blunted reward outcome value. The Dynamic Vulnerability Theory and its revised form (R-DVM) reconciles these seemingly discrepant effects by suggesting that an initial hyper-responsivity of the reward system to taste may lead to greater cue-reward learning and habituation to food reward, both of which promote overeating (Stice et al., 2011, Burger & Stice, 2011; Stice & Yokum, 2016). This account argues that the differences in hyper- and hypo-reactivity to reward occur at different times in the development of obesity, with hyporeactivity a consequence of repeated overeating, rather than an initial driver of consumption behaviour. It is unclear, however, why studies using samples with already-overweight participants have reported differences in reward responses. Irrespective of when this blunted reward response to food occurs in the development of obesity, it might be of critical importance. Blunted reward response to food delivery coupled with the increased expectation triggered by food cues may be particularly potent in driving over-consumption to compensate for experienced dissatisfaction.

Given the importance of the pattern of neural changes in response to food and food cues, both independently and in tandem, for out understanding of overweight, we sought to extend the spatial information gained from fMRI. We harnessed the superior temporal resolution of EEG to chart the time course of taste processing in overweight and normal-weight participants during anticipation and delivery of pleasant and neutral taste stimuli. In order to mimic the conditioning process leading to a heightened expected reward value, we conditioned participants to associate certain geometric shapes (cue) with the delivery of sweet, pleasant taste stimuli (squash), and other geometric shapes with a neutral taste stimulus (control). Below, we describe predictions for how overweight may modulate ERP components at cue and taste onsets (see Table 1 for a summary). We distinguish between early and late components to show that in addition to addressing the theoretical predictions of R-DVM, EEG can also help decide whether differences in the overweight group emerge from early perceptual processing of the cue or from conscious processing.

**Table 1:** Predicted effects of weight status on ERPs of interest and the corresponding observed effects in our data.

| Onset | Processing | Component | Predicted main effect of taste (sweet vs. neutral) | Predicted modulation by weight status | Observed main effect of taste | Observed valence x weight status |
| --- | --- | --- | --- | --- | --- | --- |
| Cue | Early | N1 | N1 more negative for sweet | N1 more negative for overweight | No | N1 more negative for overweight |
| Cue | Early | P2 | P2 more positive for sweet | P2 more positive for overweight | No | n.s. |
| Cue | Late | N2 | N2 more negative for neutral (fronto-central) OR N2 more negative for sweet (cento-parietal) | N2 more negative for overweight for neutral (fronto-central) or sweet (cento-parietal) | No | n.s. |
| Cue | Late | P3 | P3 more positive for sweet | P3 more positive for overweight | P3 more positive for sweet | n.s. |
| Cue | Late | SPN | SPN more negative for sweet | SPN more negative for overweight | SPN more negative for sweet | n.s. |
| Taste delivery | Early | P1 | P1 more positive for sweet | P1 less positive in overweight | No | n.s. |
| Taste delivery | Early | N1 | N1 more negative for sweet | N1 less negative in overweight | No | n.s. |
| Taste delivery | Early | P2 | P2 more positive for sweet | P2 less positive in overweight | No | n.s. |
| Taste delivery | Late | P3 | P3 more positive for sweet | P3 less positive in overweight | No | n.s. |

n.s. = not significant

### 1.1 Modulation of ERP components by obesity

#### 1.1.1 ERP responses to taste cues

There is plentiful evidence that reward expectation can modulate the N1, an early negative ERP (approximately 100ms post-cue) reflecting perceptual processing (Flores et al., 2015; Hopf et al., 2002). N1 amplitude is larger for cues signaling impending reward, compared to no-reward cues (Flores et al., 2015). This increase in N1 amplitude by reward cues is thought to reflect increased visual processing due to heightened early attention allocation to cues signalling reward (Flores et al., 2015). Given the ability of food cues to activate reward regions in a similar manner to other reward cues (Stice et al., 2011), we might expect a similar modulation of N1 by food cues. A prior study testing conditioning for sweet taste stimuli did not observe modulation of N1 for sweet relative to neutral taste (Franken et al., 2011), but their study used only normal-weight participants. If expectation effects are driven in part by increased implicit, automatic allocation of attention to food cues in overweight individuals, we would expect to see a greater enhancement of N1 by food relative to non-food cues in this group.

Food cue valence has also been demonstrated to modulate mid-latency ERPs, namely the P2 and N2. The P2 is a positive mid-latency ERP reflecting pre-conscious attention (Van Voorhis & Hillyard, 1977). Franken and colleagues (2011) observed larger P2 amplitude for cues signalling upcoming sweet relative to neutral taste stimuli. This is consistent with expecting reward from the sweet stimuli, as rewarding non-food items show a similar amplitude enhancement (Franken et al., 2011; Wang, Kleffner, Carolan, & Liotti, 2018). The effects of weight status have not yet been reported with respect to P2. A weight-group difference in P2 amplitude as a function of cue type would demonstrate a difference in pre-conscious allocation of attention that might drive the differences in reward expectation. The other mid-latency ERP of interest is the N2, a negative deflection occurring between 200 and 300ms after stimulus onset. The fronto-central subcomponent is less negative for reward cues compared to non-reward cues (Glazer et al., 2018; Novak & Foti, 2015; Pornpattananangkul & Nusslock, 2015). For food-related reward specifically, a systematic review found frontal N2 amplitude is generally more negative for neutral than palatable food cues. However, at central and posterior electrodes N2 amplitude is more negative for palatable food cues, indicating increased demand on working memory resources by these stimuli (Carbine et al., 2018). The discrepancy in N2 effects is likely due to different subcomponents being modulated by reward in different ways, given they reflect slightly different processes (Coles & Rugg, 1995). We will test the effects of food reward and weight status on the N2 to disentangle these processes and their relevance to food expected reward.

Rather than food cues triggering weight-related differences in early processes, differential responses to food cues might be primarily due to differences in higher-order, conscious processes, which modulate late-stage EPRs. The P3 is typically observed between 300 and 500ms post stimulus (Coles & Rugg, 1995), reflecting attention to salient stimuli and updating of working memory representations of stimuli (Nieuwenhuis et al., 2005). P3 amplitude is larger in response to cues indicating impending reward relative to no reward, demonstrating enhancement by reward expectation (Baines et al., 2011). Expectation of gustatory reward also modulates P3 amplitude. Cues associated with imminent delivery of a pleasant taste elicit larger P3 amplitude than those associated with neutral tastes or absence of pleasant taste delivery (Franken et al., 2011; Viemose et al., 2013). However, these studies did not look at weight status as a factor influencing this response. Studies that have tested the association of P3 amplitude and weight status are relatively few in number. They have reported larger P3 amplitude in response to pictures of palatable foods in overweight relative to normal-weight women when satiated (Nijs et al., 2010), but not when hungry (Nijs et al., 2008; but see Hill et al., 2013). We therefore predicted P3 amplitude would be larger in response to cues signalling delivery of a pleasant taste stimulus relative to neutral, and that this difference would be larger for overweight than normal-weight participants.

A particularly important marker of reward expectation is the stimulus-preceding negativity (SPN) (Brunia & Damen, 1988). The SPN reflects sustained attention to stimuli signalling imminent arrival of rewarding and punishing stimuli (Brunia et al., 2011). The SPN occurs frontocentrally (Kotani et al., 2001; Masaki et al., 2006), and is often present predominantly in the right hemisphere (Brunia & Damen, 1988; Hirao et al., 2016). Its time course is tied to the timing of the rewarding or punishing stimulus, usually occurring from 500ms before the reward stimulus onset until stimulus presentation (Fuentemilla et al., 2013; Hirao et al., 2016). No previous studies on food reward processing have examined the SPN, but in non-food reward studies the SPN is more negative when a reward is anticipated than when participants already know they will not be receiving a reward (Pornpattananangkul & Nusslock, 2015). If greater conscious expectation of reward does underlie obesity, we would expect a larger SPN amplitude in the overweight relative to normal-weight participants in response to the pleasant taste cues.

### 1.2 ERP responses to taste delivery

With respect to neural response to taste stimulus delivery, regardless of whether the processing of taste is enhanced or blunted in overweight, weight status may modulate early perceptual processing (P1 and N1). The P1 is an early positive ERP reflecting early sensory processing (Coles & Rugg, 1995; Hummel et al., 2010; Luck, 2014). It is modulated by attention, meaning a larger P1 amplitude can reflect a stronger early allocation of attention towards the stimulus (Baines et al., 2011; Luck et al., 2000; Wolz et al., 2015). For gustatory stimuli, P1 amplitude is often larger for sweet compared to neutral or less sweet stimuli (Franken et al., 2011; Ohla et al., 2012; Wilton et al., 2019). The N1 also reflects early sensory processing, in particular the intensity of taste stimuli (Tzieropoulos et al., 2013). The N1 has been observed in response to ‘electric tastes’ (stimulation of the tongue by electrical pulse) (Ohla et al., 2009, 2012), salty liquids (Mizoguchi et al., 2002), and dietary fat (Tzieropoulos et al., 2013). It must be noted that the qualities of these stimuli differ somewhat from the sweet squash stimuli used in our study, and studies using sweet tastes suggest N1 effects may not manifest (Franken et al., 2011). If overweight people indeed experience blunted taste processing, then this group should show a smaller difference in P1 and/or N1 to pleasant and neutral taste stimuli compared to normal-weight participants.

Mid-latency ERPs show modulation by both the physical qualities and reward value of taste stimuli. In addition to the aforementioned modulation by reward, P2 amplitude is also sensitive to stimulus intensity. Amplitude is larger in response to very sweet compared to weakly sweet tastes (Wilton et al., 2019). Taste stimuli with a stronger concentration may elicit more allocation of attention at the early stages of stimulus processing (Wilton et al., 2019). The sensitivity of P2 amplitude to both taste intensity and reward value suggests that if blunted reward response is driven in part by blunted stimulus processing, the difference in P2 amplitude for pleasant and neutral taste stimuli should be smaller for overweight than normal-weight participants.

Alternatively, it is possible that overweight participants show normal sensory processing of gustatory stimuli, with the modulated response occurring at conscious, deliberative processing stages, perhaps due to altered attention to gustatory stimuli. Behavioural and fMRI data regarding sensory responses in overweight participants have been contradictory. Whilst some studies show blunted response, others show no difference from normal-weight participants (Demos et al., 2012; Hardikar et al., 2017; Martinez-Cordero et al., 2015). ERP analysis will allow us to disentangle these effects. Consistent with the enhancement of P3 amplitude by rewards (Baines et al., 2011; Goldstein et al., 2006), relative to non-rewards, P3 amplitude is larger for sweet compared to neutral taste stimuli (Franken et al., 2011). If reward outcome response differences are due to a blunted experience of reward, we would expect overweight participants to show smaller differences in P3 amplitude for pleasant and neutral taste stimuli than normal-weight participants.

Evidence for differences in reward expectation and outcome as a function of weight status may be specific to food, but they could also be due to general differences in reward processing. Again, results are contradictory, with some studies demonstrating food-specific differences and others general reward deficits in overweight participants (Demos et al., 2012; Stoeckel et al., 2008). In this study we sought to address this issue by comparing reward processing to taste with pleasant and neutral pictures. We did not expect to find a difference between overweight and normal-weight people on the picture trials.

To summarise, the main aim of this study was to test the prediction of a co-occurrence of increased expectation and decreased reward outcome in overweight compared with normal-weight participants. The second aim of the study was to investigate the mechanisms underlying weight-related differences in neural responses by testing the stage(s) of processing at which weight group differences in ERPs occurred. In addition to using EEG to investigate neural responses to cues and stimuli, we also asked participants to rate the pleasantness of these tastes on every trial. This was to examine whether there was a difference between overweight and normal-weight people’s consciously-perceived enjoyment of the taste stimuli, and the relationship between this and the neural response.

## 2. Methods

### 2.1 Participants

Given the reported difference in taste and food stimulus processing between men and women (Hummel et al., 2010; Wang, Volkow, et al., 2009; Watson & Garvey, 2013), only women were recruited in order to keep the sample homogeneous. Participants were sampled from the general population using opportunity sampling. The inclusion criteria consisted of: being either normal-weight (BMI between 18.5 and 24.9) or overweight/obese (BMI of 25.0 or over); having normal or corrected-to-normal vision; age 18 years or older; and sufficient English proficiency to understand the study instructions. Exclusion criteria were: history of eating disorders (e.g. anorexia nervosa), although people with BED were not excluded; history of psychological illness (e.g. depression); history of neurological illness (e.g. epilepsy); using medication that does not allow fasting, or influences satiety, hunger, gut hormone levels, or dopaminergic activity; disease that does not allow fasting; diabetes; participating in a calorically restrictive diet during the three months prior to the experiment; being a smoker or regular recreational drugs user; having a food intolerance if this intolerance interfered with the taste stimuli; being pregnant or breastfeeding at the time of the experiment. To screen for depression, participants were asked whether they had ever been diagnosed with depression or if they thought they suffered from depression at that time. The study took place at the University of Manchester, and ethical approval was granted by the University Research Ethics Committee.

63 participants completed the entire experiment: 39 normal-weight individuals and 24 overweight individuals. Weight cut-offs were based on the National Health Service criterion for overweight (National Health Service, 2016) and on previous studies investigating reward processing in overweight people (e.g. Hendrikse et al., 2015; Nijs et al., 2008). 10 participants were excluded because they rated the neutral taste as more pleasant than the pleasant taste). One additional person was excluded because they were the only person in the sample who met the Diagnostic and Statistical Manual of Mental Disorders (DSM) criteria for binge eating disorder. Of these 11 excluded participants, nine were normal-weight and two were overweight or obese. 52 participants (22 overweight and 30 normal-weight) were retained for analysis of the behavioural data. For the EEG data analysis, one additional overweight person was excluded because of an excessive number of noisy EEG channels (see section 2.7.2).

### 2.2 Materials

#### 2.2.1 Picture selection

The picture stimuli were taken from the International Affective Picture System (IAPS; Lang, Bradley, & Cuthbert, 2008). The pictures were chosen to be either pleasant or neutral based on the ratings provided. 60 unique pictures were used for each condition. All pictures were 1024 x 768 pixels. Both picture categories were matched on content (a range of people, animals, objects, and views) and arousal ratings (Pleasant M=5.15, SD=0.42 and Neutral M=5.16, SD=0.62). The moderate arousal ratings were selected to increase the likelihood that participants attended to all pictures (Balsamo, Carlucci, Padulo, Perfetti & Fairfield, 2020).). The pleasant pictures^1^ had an average pleasantness rating of 7.76 out of nine (SD=0.50, min=7.03, max=8.74). The neutral pictures^2^ had an average pleasantness rating of 4.95 out of nine (SD=0.62, min=4.05, max=5.99), with nine being the most pleasant rating.

#### 2.2.2 Questionnaires and measures

##### 2.2.2.1 Dutch Eating Behaviour Questionnaire (DEBQ)

The DEBQ (van Strien et al., 1986) consists of three subscales measuring restrained, emotional, and external eating. This questionnaire was included because all three have been identified as potentially important moderators of the link between weight status and food reward processing, and because prior work has suggested that emotional eating is especially relevant in the context of conditioning (Hensels & Baines, 2016). Therefore, we wanted to check whether the two groups differed on these measures to eliminate them as possible confounds. The DEBQ consists of 33 items: ten external eating items, ten restrained eating items, and 13 emotional eating items. Each question was answered on a five-point Likert scale (1=never to 5=always). The answers to the questions were averaged per subscale, resulting in a separate score for every subscale ranging from one to five. Higher scores indicated higher levels of restrained, emotional, and external eating. The DEBQ has previously been shown to have excellent reliability for each of the three scales (Hensels & Baines, 2016). In the current study, the restrained and emotional eating subscales show excellent reliability (α=.864 and α=.929 respectively). The external eating subscale showed reasonably good reliability (α=.776).

##### 2.2.2.2 Binge eating disorder (BED)

BED is another potentially important moderator of weight status and food reward (Littel et al., 2012; Svaldi et al., 2010). People suffering from BED experience more neural activation in response to food cues in the OFC and the insula (Schienle et al., 2009), areas traditionally related to taste processing (Rolls, 2014). According to the DSM-V, a person suffers from BED if they report recurrent episodes of binge eating, which must be characterised by eating an amount of food in a short amount of time that is much more than an average person would consume in that same time, and by a feeling of lack of control over the food consumption. These episodes must also be associated with at least three of the following: eating a lot while not actually hungry; eating alone because of the embarrassment over the amount eaten; eating much faster than usual; eating until one is uncomfortably full; feeling guilty, depressed, or disgusted after eating. These episodes should occur once a week at a minimum for at least three months. To be diagnosed with BED, a person also needs to feel a level of stress over the binge eating. Lastly, the binge eating must not occur in tandem with extreme compensatory behaviours, such as purging (American Psychiatric Association, 2013). Only one participant in the current sample met the DSM criteria for BED, so to reduce noise in the data this person was excluded from the analysis of the behavioural and EEG data.

##### 2.2.2.3 Reward responsiveness

The reward responsiveness scale (van den Berg et al., 2010) was utilised to test whether there were differences between the normal-weight and overweight groups on general, non-food-specific reward processingThis measure of general reward sensitivity includes no items referring to reward experienced from food or drink, thus was appropriate to determine that differences in reward processing between overweight and normal-weight participants were specific to food, and that there was no generalised difference in reward sensitivity. The reward responsiveness scale consists of eight items (e.g. “If I discover something new I like, I usually continue doing it for a while”), all rated on a four-point Likert scale (1=strong disagreement to 4=strong agreement). Higher scores indicated stronger reward responsiveness. The scale has previously shown good internal consistency, with Cronbach’s alphas between 0.71 and 0.85, and good test-retest reliability (van den Berg et al., 2010). In the current study, the scale showed reasonably good reliability (α=.747).

##### 2.2.2.4 State visual analague scales (VASs)

Hunger, fullness, food craving, and thirst were tested throughout the experiment using four ten-point scales. Participants were asked was “how hungry/full/thirsty do you feel right now?” (1=not hungry/full/thirsty at all to 10=extremely hungry/full/thirsty) and “How strong is your desire to eat right now?” (1=not strong at all to 10=extremely strong). Higher scores indicated higher levels of hunger, fullness, craving, and thirst. Measurements were taken when participants entered the lab (time 1), when they were about to start the conditioning task (time 2), and after they finished the conditioning task (time 3).

### 2.3 Procedure

Before participants came into the lab, they were sent an online pre-screen (see Appendix I). If they fulfilled the inclusion criteria, they were invited to take part in the experiment. Participants’ start time was kept between 8am and 11am, to minimise changes in hunger and craving as a result of the circadian rhythm (Scheer et al., 2013). Participants were asked to have breakfast just before coming in to ensure the experiment was conducted under conditions of satiety. After providing informed consent, the participants were asked to rate their hunger, fullness, craving, and thirst levels for the first time.

#### 2.3.1 Anthropometrics

Following these ratings, participants’ height, weight, waist circumference, and hip circumference were measured. This was to confirm that participants fell into the normal-weight group (BMI between 18.5 and 24.9) or the overweight group (BMI of 25.0 or over). Waist-to-hip ratio (waist circumference divided by hip circumference) was calculated as an accurate proxy of body fat (World Health Organization, 2011), so we could confirm that, firstly, the participant was in fact normal-weight or overweight, and secondly, if they were overweight that this was due to a high body fat percentage rather than muscle mass. Participants were then asked whether they had a regular menstrual cycle, and if so, how many days their entire menstrual cycle (including non-menses phases) usually takes (e.g. Bohon, Stice, & Spoor, 2009), how many days it had been since their last menstruation started and whether they were on hormonal contraceptives. This was done because menstrual cycle phase has been found to affect reward responsiveness (Dreher et al., 2007). By measuring this we could therefore confirm that both groups were in similar phases of their menstrual cycle at the time of testing.

#### 2.3.2 Taste stimulus selection

Two squashes, orange and apple/blackcurrant, were used in the pleasant taste condition to keep the pleasant taste trials variable, and thereby reduce the rate of sensory-specific satiety (Havermans et al., 2009). The concentrations of squash to be used in the conditioning task were tailored to each participant individually. This was done within a fixed range by measuring taste pleasantness and intensity ratings of four different concentrations of both squash flavours. The four concentrations were 20 ml, 10 ml, 5 ml, or 3 ml of squash per 40 ml of water. Prior pilot testing ensured these concentrations provided a spread in taste intensity, so we could cater to a wide range of sweetness and intensity preferences.

The four concentrations of each squash were presented five times in randomised order which was identical for all participants. Participants rated each stimulus on pleasantness and intensity on a scale from one (“not pleasant at all”/”not intense at all”) to nine (“very pleasant”/”very intense”). Only the ‘one’ and ‘nine’ markers on the scales were labelled. Participants were instructed on how to use the scales ahead of the task.

The most pleasantly-rated concentration for each squash was chosen for use in the conditioning task. One-variable chi-square tests analysing pleasantness ratings collected during the taste stimulus selection task (see Table 2) showed participants most often rated the strongest concentration (i.e. 20 ml of squash with 40 ml of water) as the most pleasant concentration for both orange, χ^2^ (3)=46.77, p<.001 and apple/blackcurrant squash, χ^2^ (3)=50.00, p<.001. A 2×2 chi-square test of pleasantness ratings showed the concentration rated most pleasant did not differ according to weight status for either orange, χ^2^ (3)=2.56, p=.464, Cramer’s V=.222 or apple/blackcurrant squash, χ^2^ (3)=3.32, p=.345, Cramer’s V=.253.

**Table 2.** Pleasantness and intensity ratings (on a scale from one to nine, nine being the most pleasant/intense) for the four piloted concentrations of orange squash and apple/blackcurrant squash. Each concentration signifies the amount of squash per 40 ml of water. For each participant individually, the concentrations of orange and apple/blackcurrant squash that were rated highest on pleasantness were chosen to be used in the conditioning task as pleasant tastes. This was most often the concentration with 20 ml of squash per 40 ml of water.

|  | **Pleasantness** |  |  | **Intensity** |  |  |
| --- | --- | --- | --- | --- | --- | --- |
|  | Total | Overweight | Normal-weight | Total | Overweight | Normal-weight |
|  | N=52 | N=22 | N=30 | N=52 | N=22 | N=20 |
| **Orange squash** |  |  |  |  |  |  |
| *3 ml* | 3.68 (1.47) | 3.85 (1.61) | 3.55 (1.38) | 2.29 (1.22) | 2.34 (1.34) | 2.25 (1.14) |
| *5 ml* | 4.47 (1.35) | 4.75 (1.21) | 4.27 (1.42) | 3.01 (1.14) | 3.26 (1.26) | 2.83 (1.02) |
| *10 ml* | 5.23 (1.16) | 5.23 (0.89) | 5.22 (1.34) | 4.57 (1.34) | 4.86 (1.23) | 4.35 (1.39) |
| *20 ml* | 5.53 (1.35) | 5.75 (1.34) | 5.23 (1.33) | 5.99 (1.55) | 6.26 (1.46) | 5.79 (1.60) |
| **Apple/ black-currant squash** |  |  |  |  |  |  |
| *3 ml* | 3.78 (1.44) | 3.92 (1.62) | 3.68 (1.32) | 2.44 (1.15) | 2.51 (1.27) | 2.38 (1.07) |
| *5 ml* | 4.54 (1.29) | 4.46 (1.40) | 4.59 (1.22) | 3.26 (1.15) | 3.22 (1.13) | 3.29 (1.19) |
| *10 ml* | 5.29 (1.21) | 5.40 (1.11) | 5.21 (1.29) | 4.46 (1.28) | 4.49 (1.31) | 4.44 (1.29) |
| *20 ml* | 5.62 (1.41) | 5.54 (1.28) | 5.68 (1.52) | 5.94 (1.59) | 5.86 (1.68) | 6.00 (1.56) |

*Notes.* Reported data are M (SD).

The neutral stimulus was distilled water with 2 mM NaHCO_3_ and 15 mM KCl. This typical control stimulus in neuroimaging studies of taste mimics the characteristics of saliva, and is thus generally considered to be chemosensorily neutral (Franken et al., 2011; Hird et al., 2017; see Baines, Hensels & Talmi, 2020 for discussion). The neutral stimulus was the same for all participants, and was not administered prior to the conditioning task. Upon completion of the taste stimulus selection task, participants were given a ten-minute break and urged to eat some food during this break if they were still feeling “peckish”.

#### 2.3.3 Conditioning task

After their ten-minute break, participants were set up with EEG electrodes. Once setup was complete, participants again rated their hunger, fullness, craving, and thirst levels before commencing the conditioning task. This was implemented in MATLAB R2007a using the Image Processing and the Data Acquisition Toolboxes (The MathWorks Inc., 2007). First participants were taught to associate geometric shapes with pleasant and neutral taste and pictures stimuli. Green and orange squares and triangles were associated with pleasant and neutral tastes and pictures. The colour of the cue always indicated the stimulus type (e.g. green for tastes and orange for pictures). The shape of the cue always indicated the stimulus valence (e.g. circle for pleasant and triangle for neutral). For each participant, the same shape was always associated with the same kind of stimulus. So for instance the green circle always signalled arrival of a pleasant taste, the green triangle a neutral taste, the orange circle a pleasant picture and the orange triangle a neutral picture. Cue-stimulus assignment was counterbalanced across participants.

During the conditioning task, participants completed 60 trials for each of the four conditions, making a total of 240 trials. The order of trials was randomised for each participant, and the trials were presented in four blocks of 60 trials, with a break between blocks. The progression of events within each trial (see Figure 1) was modelled after Franken et al. (2011) and Hird, El-Deredy, Jones, and Talmi (2017). Each trial consisted of the following: a fixation cross (jittered between 500-750ms), followed by presentation of the cue (750ms). In the taste condition a screen reading ‘taste’ followed, during which the taste was delivered (1000ms) and held in the mouth (1000ms). This was followed by a screen reading ‘swallow’, during which the participant was instructed to swallow the stimulus (2000ms). In the picture condition the presentation of a picture followed the cue (2000ms). Then, participants were asked to rate the pleasantness, intensity, and comparative pleasantness of the presented stimulus on three successive Likert scales. At the end of each taste trial, the participants received a wash of the neutral stimulus to cleanse the palate (1250ms). A jittered inter-trial interval (500-1500ms) followed all trials. All taste stimuli were administered at 10 ml/min. During the task, participants were asked to wear earplugs to mask the sound of the syringes administering the stimuli, to ensure they did not know exactly when the stimulus was administered. The task lasted approximately 70 minutes. After completing the conditioning task, participants were asked to once again rate their current hunger, fullness, craving, and thirst levels and they were asked to fill in the DEBQ, the reward responsiveness questionnaire, and the BED questionnaire. See Figure 2 for a schematic overview of the entire timeline of the experimental procedure.

**Figure 1.**
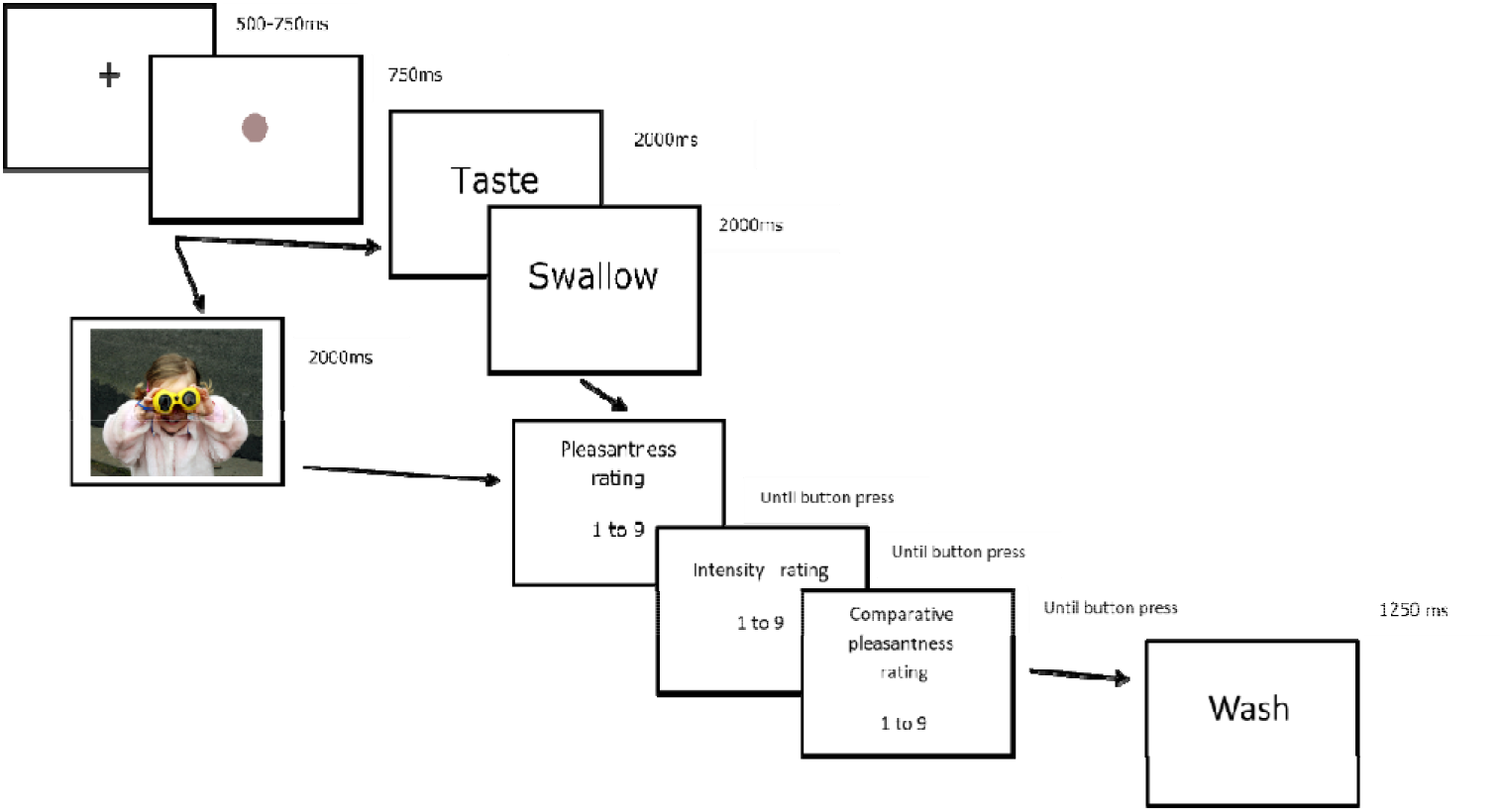
Time course of a trial of the conditioning task. A fixation cross (500-750ms) was presented, followed by a central cue (750ms, geometric shape). Next, a taste (see trial progression on the right) or picture (see trial progression on the left) stimulus was presented (2000ms). A 2000ms interval (with the instruction to swallow on taste trials) was followed by three consecutive ratings (pleasantness, intensity, comparative pleasantness). Participants could take as long as they wanted to give each rating. A schematic representation of the questions is given in this figure. For the full phrasing and rating scales of the questions, see section 2.6. After the taste trials only, participants received a wash to cleanse their palate. All trials were followed by a jittered inter-trial interval (500ms-1500ms).

**Figure 2.**
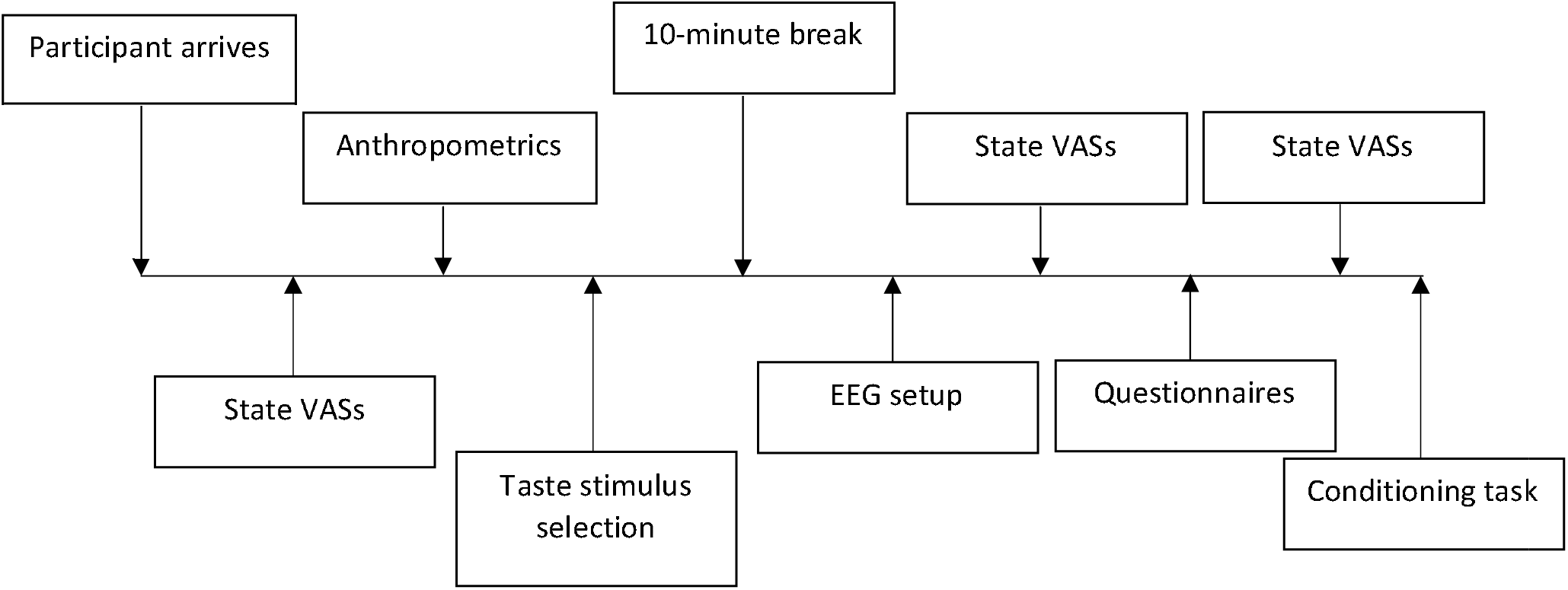
Schematic overview of the experimental timeline, in chronological order from left to right. The state VASs included ratings of hunger, fullness, craving, and thirst. The questionnaires included the Dutch Eating Behaviour Questionnaire, the Reward Responsiveness questionnaire, and a questionnaire diagnosing Binge Eating Disorder. See section 2.3 for a detailed description of each element of the procedure.

### 2.4 Taste delivery setup

The taste stimuli were delivered with the Harvard Apparatus Model 33 using four 50ml plastic syringes from Fisher Scientific. Two syringes were filled with the neutral stimulus, and one syringe was filled with each squash flavour. The pumps were placed outside the testing booth, with taste stimuli were delivered through silicone tubes with a 2mm bore and a 0.5mm wall from Altec Products Ltd. The tubes were connected to the syringes using luer adaptors from Cole-Parmer, and taped to a chin rest in order to keep the tubes and the participants’ heads in place and to minimise muscle artefacts. MATLAB R2007a (The MathWorks Inc., 2007) was used to control the pumps, present the visual stimuli, and collect the data.

### 2.5 EEG data collection

The conditioning task was completed in a sound-attenuated, electromagnetically-shielded booth with the lights dimmed. The EEG data was continuously recorded at 512 Hz using a 64-electrode Active-Two amplifier system with a ground electrode (BioSemi) and four external electrodes to measure vertical and horizontal electro-oculograms (EOGs). The electrodes were affixed to the scalp using an elasticated cap. External electrodes were positioned above and below the right eye, to the right of the right eye, and to the left of the left eye. This was done to detect blinks and eye-movements. ActiView was used for data acquisition (BioSemi). Impedance was kept below 30 KΩ.

### 2.6 Design

This study was a 2 x 2 x 2 mixed design. The between-subjects independent variable was weight status, with two levels: normal-weight and overweight/obese. The first within-subjects variable was stimulus valence, with two levels: pleasant and neutral. The second within-subjects variable was stimulus type, with two levels: taste and picture. Owing to their different modalities, responses to taste and picture were not directly compared to each other. Data treated as two separate 2 (weight) x 2 (valence) designs.

There were three behavioural dependent variables: pleasantness, intensity, and comparative pleasantness ratings. All three variables were rated on a Likert scale from one to nine. For the pleasantness and intensity ratings, participants were asked: “how pleasant/intense did you find this stimulus? Please rate it on a scale from 1 to 9” (1=not pleasant/intense at all, 9=very pleasant/intense). Ahead of the task, participants had been instructed to use the middle of scale (five) as neutral for pleasantness. The third Likert scale, labelled here as comparative pleasantness, was used to gather a self-report of how the expected outcome compared to the actual outcome. The participants were presented with the question “how did the actual pleasantness of this stimulus compare to the pleasantness you expected? Was it worse, better, or the same?” (1=much worse, 5=as expected, 9=much better). It was explained to the participants that in this instance, a rating of one indicated that the stimulus (i.e. the taste or the picture) was much worse than they expected it to be when they saw the cue, a rating of nine meant that it was much better, and a rating of five meant that it was as pleasant as they had expected it to be when they saw the cue.

### 2.7 Data preprocessing and analysis 2.7.1 Behavioural data

The behavioural data were preprocessed and analysed in SPSS 22 (IBM Corp, 2013). Manipulation checks were performed using one-sample t-tests, paired-samples t-tests, and chi-square tests. Group and valence differences in the behavioural data were investigated using six mixed 2 x 2 (weight status by valence) ANOVAs.

#### 2.7.2 EEG data preprocessing

The EEG data were preprocessed using SPM12 (Ashburner et al., 2016) in MATLAB R2015a (The MathWorks Inc., 2015). Two epochs of EEG data were analysed. Firstly, the 750ms time window in which the cue is presented was analysed to look at anticipation of the taste and picture stimuli; the ERPs in this window will be referred to as cue-elicited ERPs. Secondly, the 1000ms interval following the start of the taste or picture presentation was analysed to investigate ERPs in response to the target stimuli; these ERPs will be referred to as the target-elicited ERPs. After conversion from Biosemi to SPM format, the data were re-referenced to the average of all electrodes except for P2 and PO4, as these electrodes were very noisy across participants so removed from analysis. Butterworth highpass and lowpass filters of 0.1 and 20 Hz respectively were applied. The data were downsampled to 200 Hz. After this, topography-based artefact detection based on the vertical EOG channels was applied to identify blinks. To remove the blinks, epochs were created around the blinks (spanning 500ms before to 500ms after the peak of the blink). Then, for each participant, the first six components in these epochs were inspected. Subsequently, the number of components up to and including the component showing the blinks was removed from the data. On average, two components were removed per participant (min=1, max=5). Then, the data were epoched according to the intervals of interest (-100ms to 750ms for cues, -850ms to 1000ms for targets). For the cue-elicited ERPs, a baseline correction was applied using -100ms to 0ms of the epoch as the baseline. For the target-elicited ERPs, the epoch was baseline-corrected for the same 100ms leading up to the cue, that is, the period from 850ms to 750ms before the target stimulus. This was done because the target delivery immediately followed the cue, and using the 100ms before target onset would have overlapped with the end of the cue presentation and therefore skewed the baseline. Following baseline correction, trials where the signal in any EEG channel exceeded +/-150 μV were excluded. Trials were averaged by condition using robust averaging, after which another 20 Hz lowpass filter was applied, to remove any high-frequency noise introduced by the averaging procedure. After artefact detection, the final participant sample (N=51) had a mean of 54 trials for each condition in the cue-elicited epoch (min=35, max=60) and 50 per condition in the target-elicited epoch (min=29, max=60). On average, two channels per participant were marked as bad (min=2, max=6) for both epochs, including P2 and the PO4, which were marked as bad for everyone. One participant was excluded because they had more than 10 bad channels: 9 in the cue-elicited epoch, and 22 in the target-elicited epoch. A channel was marked as bad if more than 20 percent of all trials on that channel exceeded +/-150 μV. No additional participants had to be excluded because their data had too few usable trials or too many bad channels. For the exploratory whole-brain analysis that was carried out in SPM, the EEG data were converted to .nifti images, which were smoothed using a 6 x 6 x 25 full-width at half maximum of the Gaussian kernel.

#### 2.7.3 EEG data analysis

ERPs of interest were identified based on prior literature. Guided by this, time windows for analysis were defined using visual inspection of grand mean waveforms and topographies. A grand mean topography is given in figures 5-10 for each component, which is averaged across all valence and weight status conditions. Planned analysis used a mixed ANOVA with the factors of valence, weight status, electrode site, and hemisphere. Where either electrode site or hemisphere interacted with valence and weight, follow-up ANOVAS were conducted for the significant electrode site or hemisphere. Where interactions were followed up with simple effects, a Bonferroni correction was used. Greenhouse-Geisser statistics were reported where the assumption of sphericity was violated. In the cue-elicited epoch, we analysed the N1, the P2, the N2, the P3, and the SPN (see section 3.3.1). In the target-elicited epoch, we analysed the P1, the N1, the P2, the N2, and the P3 (see section 3.3.2).

**Figure 3.**
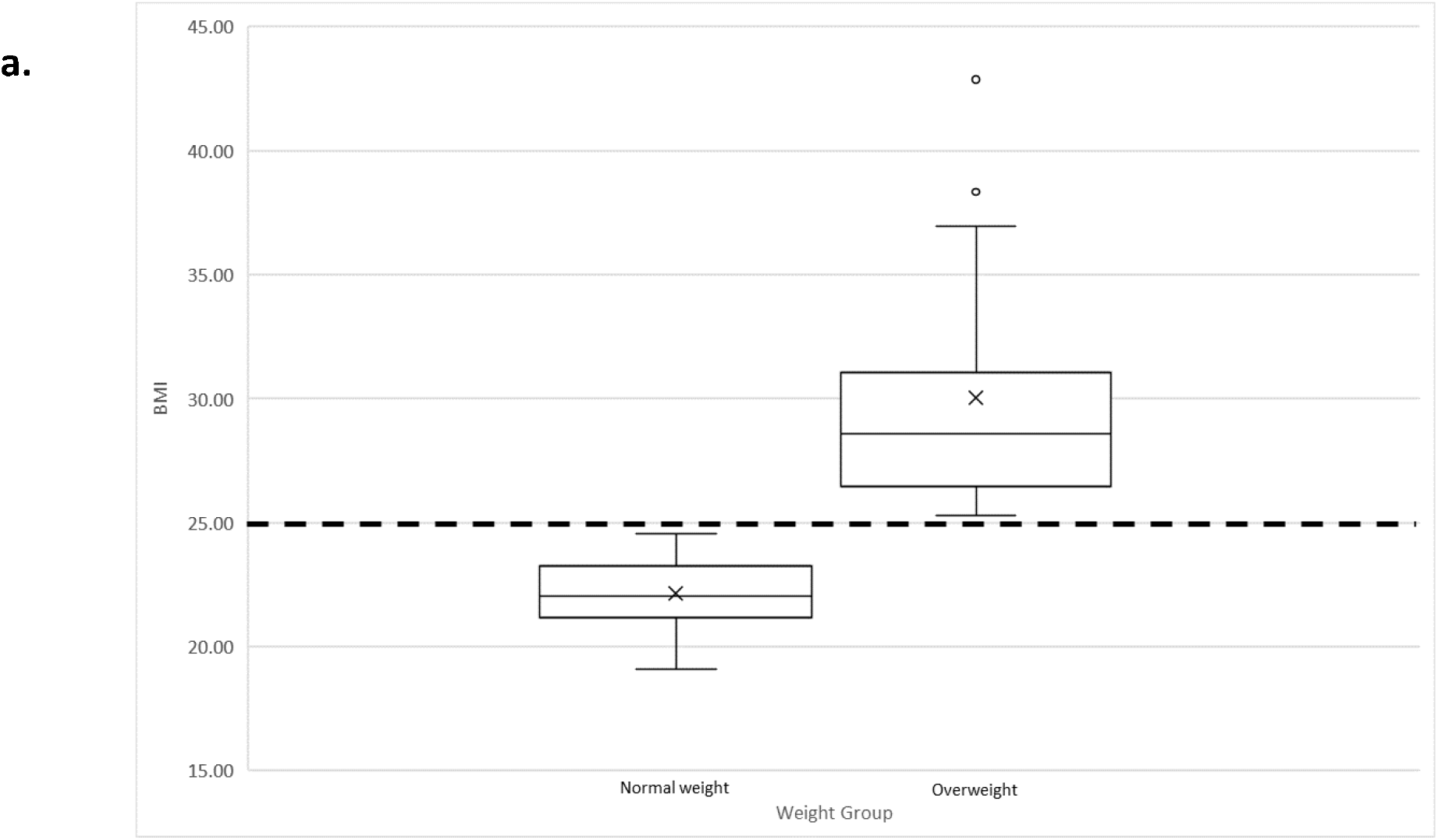
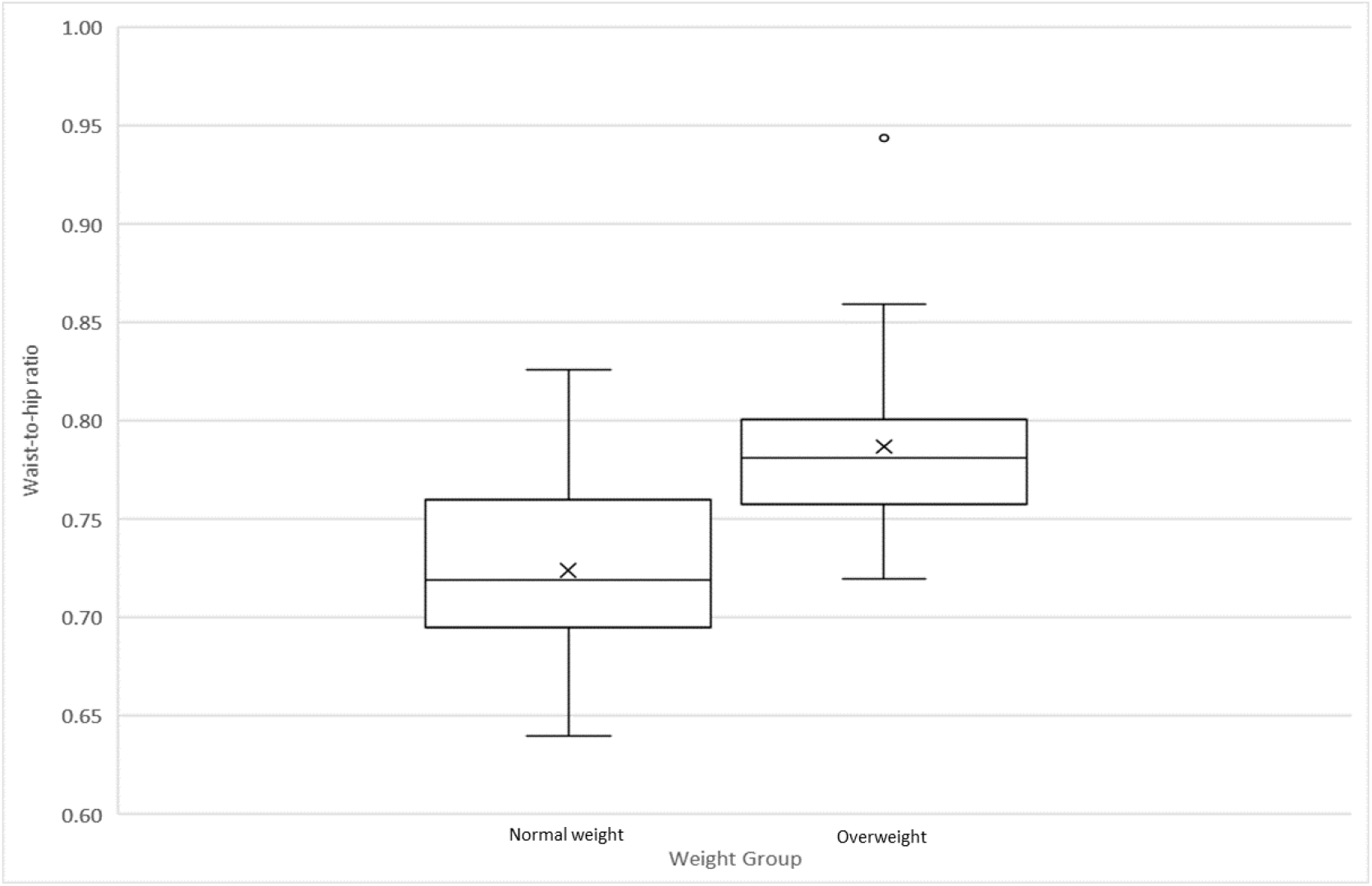
Boxplots showing the distribution of BMIs (a.) and waist-to-hip ratio (b.) for the normal-weight and overweight group. Dashed line showed the BMI cut-off used for weight-group classification.

**Figure 4.**
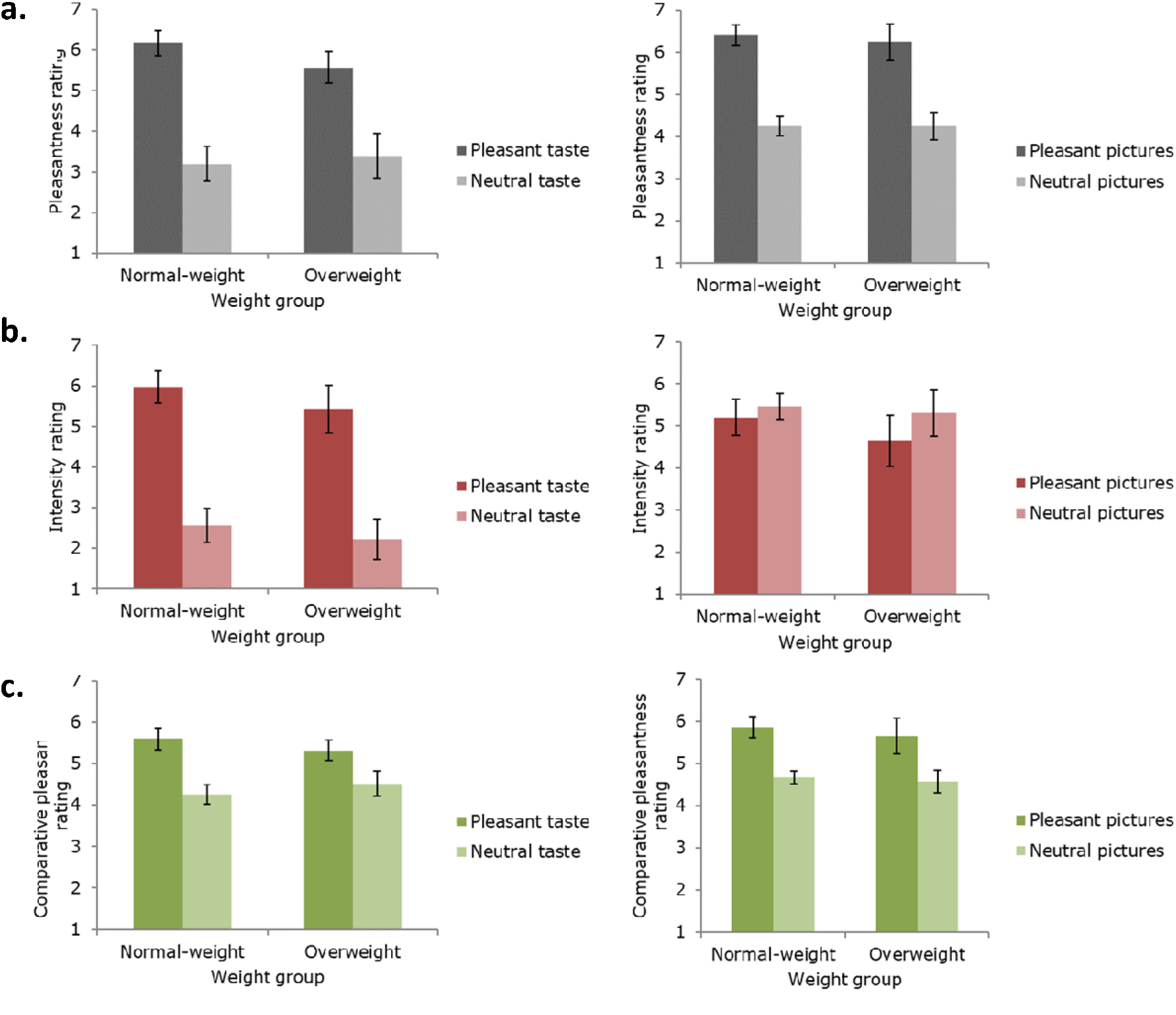
Descriptive statistics of the taste (left panels) and picture (right panels) stimulus ratings for pleasantness, intensity, and comparative pleasantness as a function of weight status. Error bars represent 95% confidence intervals. a. Pleasantness ratings. b. Intensity ratings. c. Comparative pleasantness ratings.

**Figure 5.**
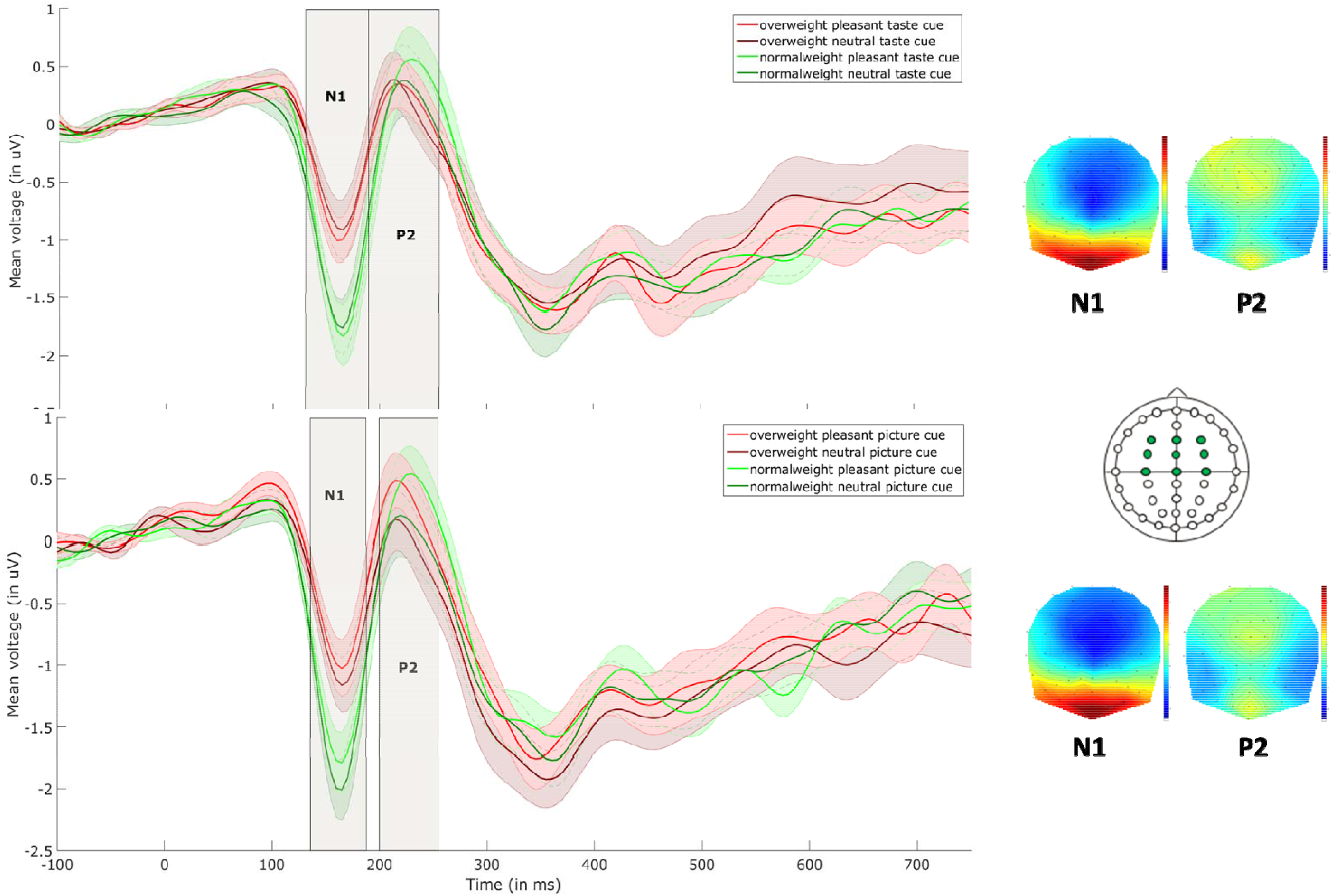
Cue-elicited ERPs. Figure shows data for taste (upper panel) and picture (lower panel) cues at electrodes F3/z/4, FC3/z/4, and C3/z/4 (marked in green on the headmap). From left to right, the topographies show scalp activity at 165ms (showing the N1, analysed between 130 and 190ms), and at 225ms (showing the P2, analysed between 190 and 265ms). The shaded areas around the waveforms represent the standard error.

**Figure 6.**
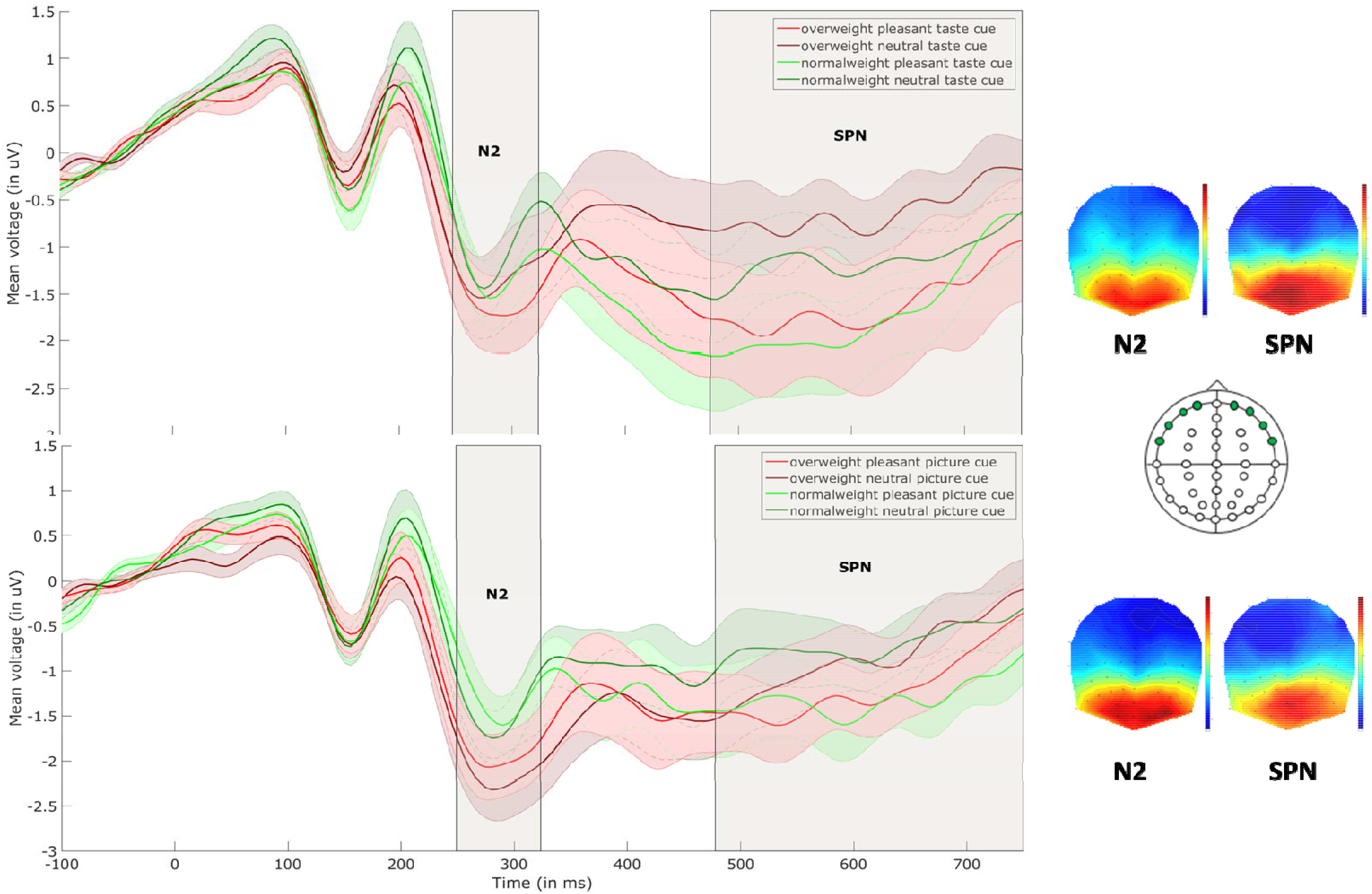
Late stage cue-elicited ERPs. Figure shows taste (upper panel) and picture (lower panel) cues at electrodes Fp1/2, AF7/8, F7/8, and FT7/8 (marked in green on the headmap).

**Figure 7.**
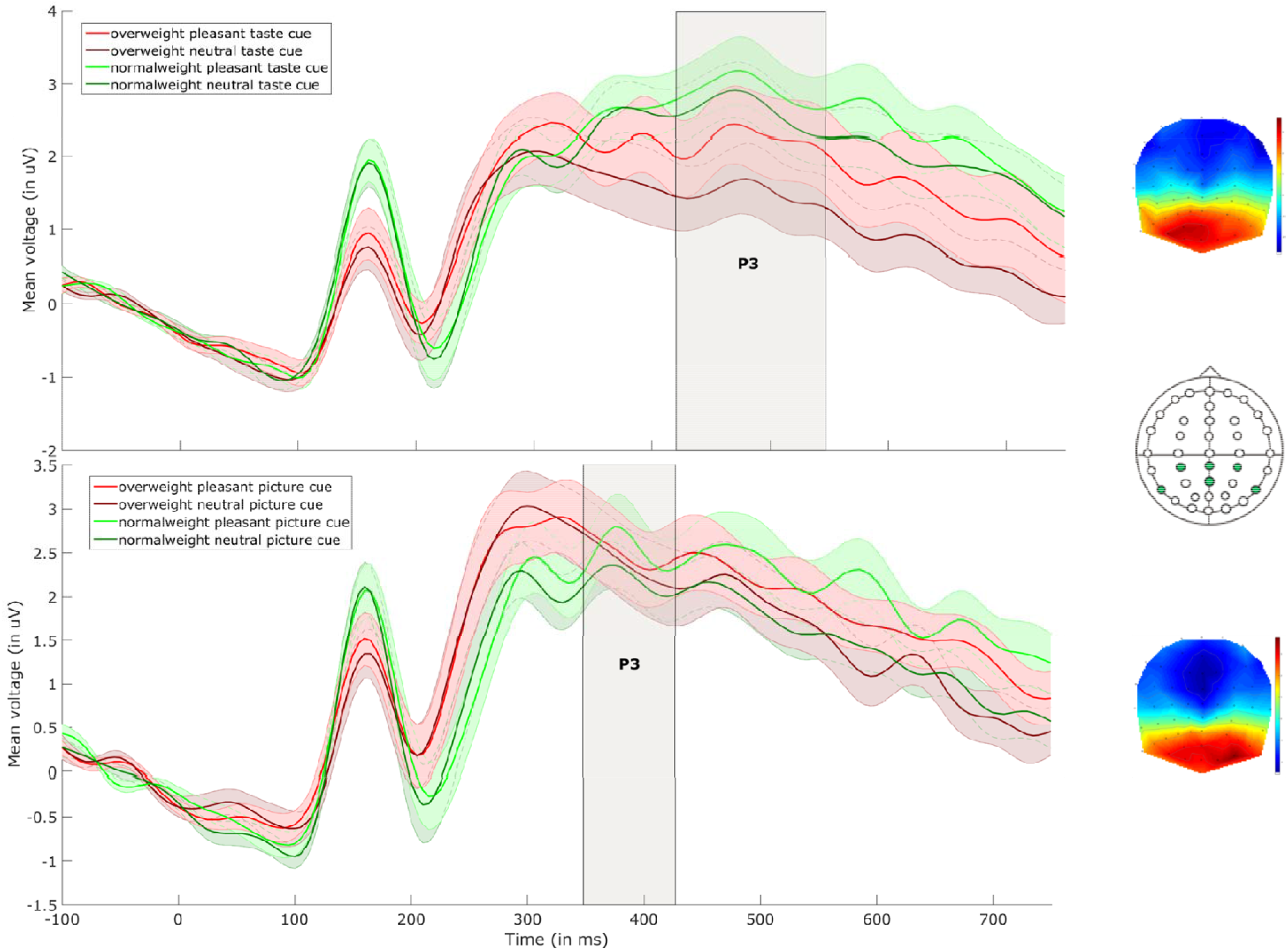
Cue-elicited P3. Figure shows taste (upper panel) and picture (lower panel) trials at electrodes P3/z/4 and PO7/z/8 (marked in green on the headmap). Topography for the taste trials shows scalp activity at 450ms (showing the P3, analysed between 420 and 550ms). Topography for the picture trials shows scalp activity at 375ms (showing the P3, analysed between 350 and 420ms). The shaded areas around the waveforms represent the standard error.

**Figure 8.**
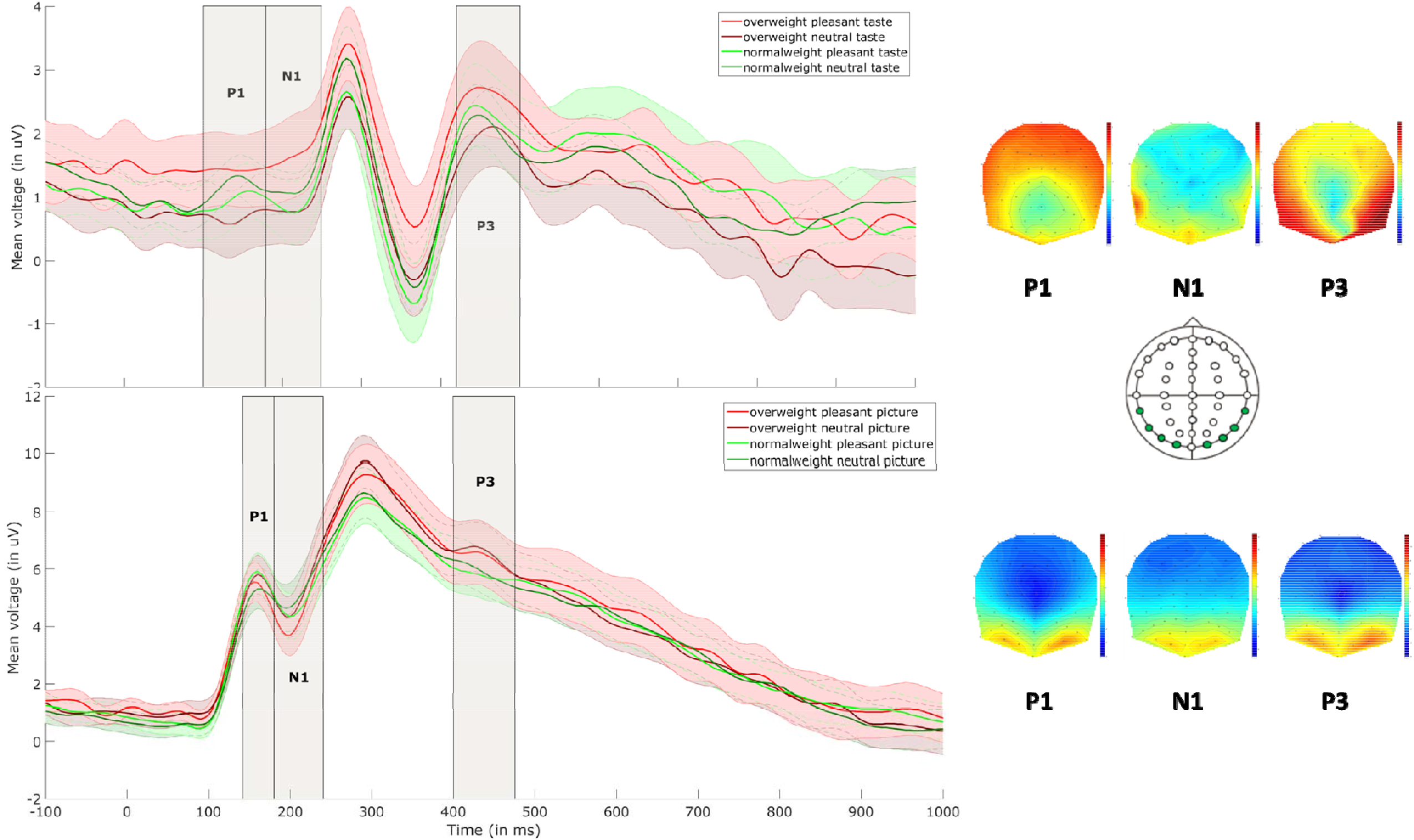
Target-elicited ERPs. Figure shows data for taste (upper panel) and picture (lower panel) trials at electodes TP7/8, P9/10, PO7/8, and O1/2 (marked in green on the headmap). From left to right, the taste topographies show scalp activity at 140ms (showing the P1, analysed between 100 and 180ms), at 220ms (showing the N1, analysed between 180 and 250ms), and at 450ms (showing the P3, analysed between 420 and 500ms). The picture topographies show scalp activity at 165ms (showing the P1, analysed between 150 and 190ms), at 200ms (showing the N1, analysed between 190 and 240ms), and at 420ms (showing the P3, analysed between 400 and 480ms). The shaded areas around the waveforms represent the standard error

**Figure 9.**
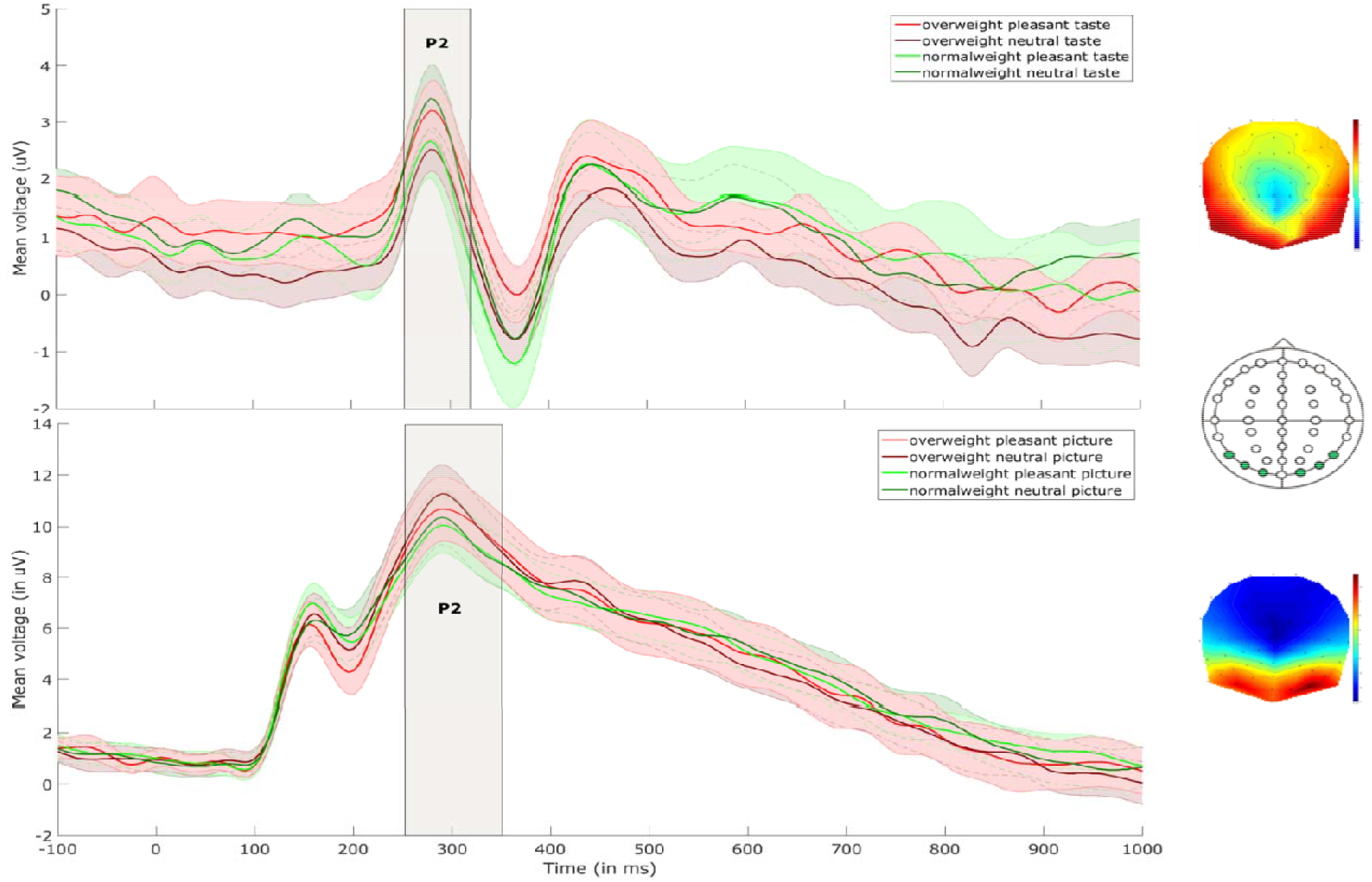
Target-elicited P2. Figure shows data for taste (upper panel) and picture (lower panel) trials, averaged over electrodes P9/10, PO7/8, and O1/2 (marked in green on the headmap). Taste topography shows scalp activity at 290ms (showing the P2, analysed between 260 and 320ms). The shaded areas around the waveforms represent the standard error. Picture topography shows scalp activity at 300ms (showing the P2, analysed between 260 and 350ms).

**Figure 10.**
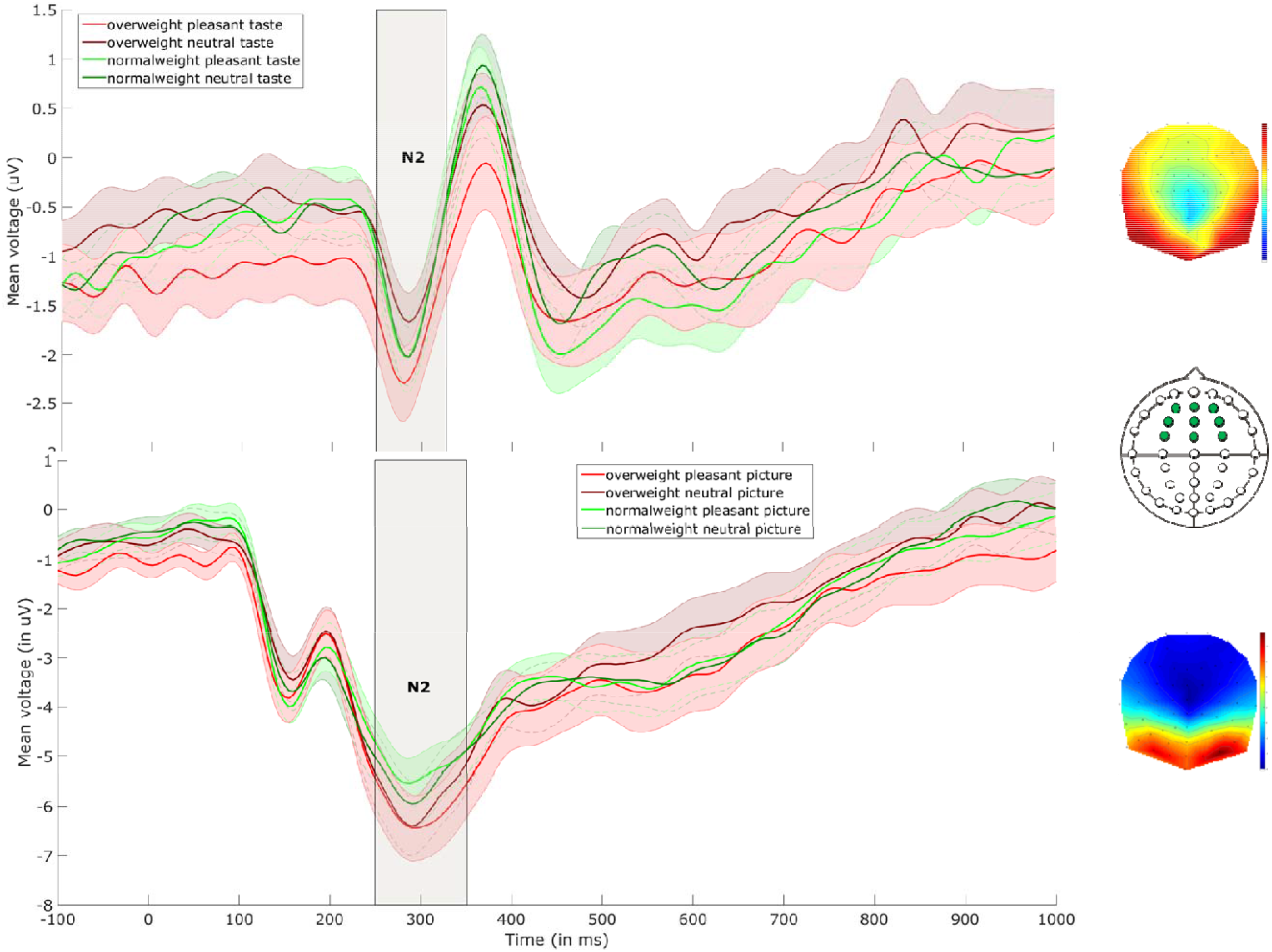
Target-elicited N2. Figure shows data for taste (upper panel) and picture (lower panel) data averaged over electrodes AF3/z/4, F1/z/2, and FC1/z/2 (marked in green on the headmap). Topographies show scalp activity at 285ms (taste) and 300ms (pictures) (analysed between 250 and 320ms). The shaded areas around the waveforms represent the standard error.

As this was an exploratory study in a field with relatively few similar previous studies, we also ran a post-hoc whole-brain analysis on both epochs, using an uncorrected voxel significance level of p<.01, and a cluster threshold of k≥200, to explore any significant clusters beyond the electrodes that were selected a priori and analysed above. For this analysis the EEG data were converted to .nifti images, which were smoothed using a 6 x 6 x 25 full-width at half maximum of the Gaussian kernel. For this exploratory analysis, clusters significant at an uncorrected cluster significance threshold of p<.05 are reported.

## 3. Results

### 3.1 Sample characteristics

As expected, the overweight group had a significantly higher BMI and a higher waist-to-hip ratio than the normal-weight group (Figure 3). The overweight group were significantly more hungry than the normal-weight group at time 1,just after they entered the lab, but not at time 2 (i.e. right before the beginning of the conditioning task) or 3 (i.e. immediately after the completion of the conditioning task). Overweight participants also reported craving food more than normal-weight participants at time 3 but not at time 1 or time 2. No group differences were found on thirst and fullness. Descriptive statistics for the sample can be found in Table 3.

**Table 3.**
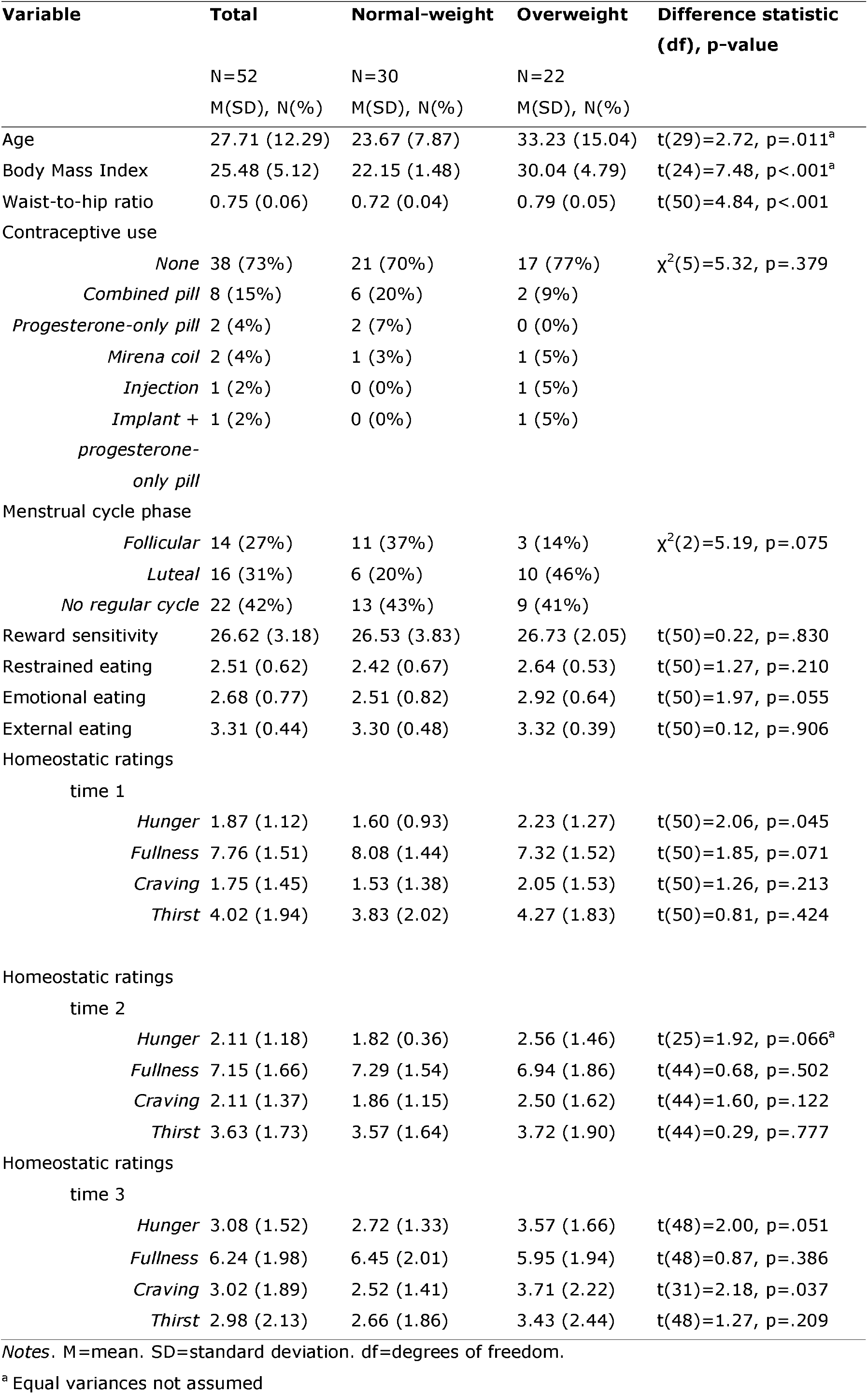
Overview of the sample demographics, split by weight status (normal-weight and overweight/obese). Difference statistics are chi-square for percentages, and t-values for means.

Unexpectedly, the overweight group was also significantly older than the normal-weight group. To check that these group differences did not skew the results, we ran all analyses with and without age as a covariate. There were no differences, so we report the results without the age covariate.

### 3.2 Behavioural data

#### 3.2.1 Manipulation check: Evidence of learning

We expected that as time went on, participants became able to predict more accurately what the neutral stimulus would taste like, as this was the same on every neutral taste trial. Comparative pleasantness of this stimulus should therefore have become increasingly close to ‘as expected’ as the task progressed. This was indeed evident in a significant positive correlation between comparative pleasantness ratings and trial order (r=.539, p<.001), indicating participants were learning to predict upcoming stimuli from the cues, even though the pleasant taste stimulus alternated between one of two squashes and was therefore less easy to predict.

#### 3.2.1 Taste pleasantness ratings

Pleasantness, intensity, and comparative pleasantness ratings for taste stimuli can be found in Figure 4. A 2 (weight status) x2 (valence) mixed ANOVA was conducted to investigate how pleasantness differed according to valence and weight status. There was a main effect of valence, with pleasant stimuli rated as more pleasant than neutral stimuli, F(1, 50)=186.40, p<.001, η^2^_p_=.788. There was no main effect of weight status, F(1, 27)=0.13, p=.725, η2_p_=.005. There was a significant interaction between valence and weight status, F(1, 50)=4.59, p=.037, η^2^_p_=.084. This interaction was due to the difference in pleasantness ratings for pleasant and neutral stimuli being much larger for the normal-weight than overweight group (see Figure 4a). Four planned contrasts were then conducted. Using two paired-samples t-tests, one for each weight group, we found that both the normal-weight, t(29)=12.84, p<.001, d=3.01, and overweight groups, t(21)=7.08, p<.001, d=2.08, rated the pleasant taste as more pleasant than the neutral taste. Two additional independent t-tests, comparing the two weight groups on the pleasant and neutral stimuli, revealed that while there was no significant difference between the two groups on pleasantness ratings for the neutral stimulus, t(50)=0.60, p=.548, d=0.17, the normal-weight group rated the pleasant stimulus as significantly more pleasant than the overweight group, t(50)=2.56, p=.014, d=0.72 (see Figure 4a). This demonstrates successful pleasantness manipulation. Furthermore, the weight group differences suggest pleasant stimuli were reduced in reward value in the overweight group, in accordance with our hypothesis.

#### 3.2.2 Taste intensity ratings

Another 2 (weight status) x2 (valence) mixed ANOVA was conducted to investigate how intensity of the two taste types differed, and whether this was dependent on weight status. There was a main effect of valence, showing pleasant stimuli were rated as more intense than neutral stimuli (see Figure 4b), F(1, 50)=249.65, p<.001, η^2^_p_=.833. There was no main effect of weight status, F(1, 50)=3.33, p=.074, η^2^_p_=.062, and no significant interaction, F(1, 50)=0.24, p=.625, η^2^_p_=.005.

#### 3.2.3 Taste comparative pleasantness ratings

Another 2 (weight status) x2 (valence) mixed ANOVA was conducted to investigate how comparative pleasantness of the stimuli (i.e. whether the taste was more or less pleasant than expected when encountering the cue) differed according to valence and weight status (see Figure 3c). There was a main effect of valence, with pleasant stimuli were more pleasant in comparison to what was expected than neutral stimuli, F(1, 50)=66.98, p<.001, η^2^_p_=.573. There was no main effect of weight status, F(1, 50)=0.01, p=.926, η^2^_p_<.001. There was a significant interaction, F(1, 50)=4.31, p=.043, η^2^_p_=.079. As can be seen in Figure 4c, the difference between comparative pleasantness ratings of pleasant and neutral stimuli was larger in the normal-weight than overweight group. Two planned contrasts were then conducted. Using two paired-samples t-tests, we found the pleasant stimulus was rated as more pleasant in comparison to what was expected than the neutral stimulus for both the normal-weight, t(29)=8.03, p<.001, d=2.00, and the overweight groups, t(21)=3.92, p=.001, d=1.27. Two one-sample t-tests were also run, using p=.025 significance threshold to correct for multiple comparisons, which compared comparative pleasantness ratings for both types of stimuli to the ‘as expected’ point on the scale (i.e. 5, being the halfway point between 1 and 9). These t-tests showed that pleasant stimuli were rated as significantly more pleasant than expected, t(51)=5.28, p<.001, d=0.73, whereas neutral stimuli were rated as less pleasant than expected, t(51)=6.81, p<.001, d=0.94.

#### 3.2.4 Picture ratings

We predicted that plesant pictures would be rated as more liked than neutral pictures, but that the two groups would not show a difference. Intensity and comparative pleasantness were not expected to differ as a function of valence or weight status. For pleasantness ratings (see Figure 4a), this revealed a main effect of valence, showing that pleasant pictures were rated as more pleasant than neutral pictures, F(1, 50)=363.02, p<.001, η^2^_p_=.879. There was no main effect of weight status on picture pleasantness, F(1, 50)=0.24, p=.626, η^2^_p_=.005, and no interaction, F(1, 50)=0.67, p=.419, η^2^_p_=.013.

For intensity ratings (see Figure 4b), the ANOVA revealed a main effect of valence. Neutral pictures were perceived as more intense than the pleasant pictures, F(1, 50)=11.45, p=.001, η^2^_p_=.186. There was no main effect of weight status, F(1, 50)=1.48, p=.230, η^2^_p_=.029, and no interaction, F(1, 50)=1.98, p=.166, η^2^_p_=.038.

For comparative pleasantness ratings (see Figure 4c), there was a main effect of valence, showing pleasant pictures were rated as more pleasant compared to what was expected than neutral pictures, F(1, 50)=93.65, p<.001, η^2^_p_=.652. Two one-sample t-tests were run using p=0.25 statistical significance threshold to compare comparative pleasantness ratings for both types of pictures to the ‘as expected’ point (i.e. 5, being the halfway point between 1 and 9). These t-tests showed that pleasant pictures were rated as significantly more pleasant than expected, t(51)=7.00, p<.001, d=0.97, whereas neutral pictures were rated as significantly less pleasant than expected, t(51)=4.63, p<.001, d=0.72. There was no main effect of weight status, F(1, 50)=1.01, p=.320, η^2^_p_=.020, and no interaction, F(1, 50)=0.23, p=.637, η^2^_p_=.004.

### 3.3 ERP analysis

#### 3.3.1 Cue-elicited ERPs

##### 3.3.1.1 N1: Effect of weight status on early processing

The N1 in response to the taste cues was analysed at electrodes F3/z/4, FC3/z/4, and C3/z/4 electrodes between 130 and 190ms after cue onset (see Figure 5). We found no main effect of valence, F(1, 49)=0.28, p=.601, η^2^_p_=.006. However, there was a main effect of weight status, F(1, 49)=6.40, p=.015, η^2^_p_=.116. The N1 was more negative for the normal-weight (M=-1.28 μV, SD=0.93) compared to overweight group (M=-0.59 μV, SD=0.93). There was no significant interaction between valence and weight status, F(1, 49)=0.04, p=.848, η^2^_p_=.001. For the picture cues, the N1 was analysed over the same electrodes, between 140 and 180ms (see Figure 5). There was no main effect of valence, F(1, 49)=1.59, p=.213, η^2^_p_=.032. However, there was a significant main effect of weight status, F(1, 49)=6.98, p=.011, η^2^_p_=.125. The normal-weight group (M=-1.59 μV, SD=0.96) had a significantly more negative N1 than the overweight group (M=- 0.86 μV, SD=0.96). There was no significant interaction between valence and weight status, F(1, 49)=0.03, p=.865, η^2^_p_=.001.

##### 3.3.1.2 P2: Effect of weight status on pre-conscious attention

The P2 in response to the taste cues was analysed at electrodes F3/z/4, FC3/z/4, and C3/z/4, between 190 and 265ms after cue onset (see Figure 5). There was no main effect of valence, F(1, 49)=0.82, p=.370, η^2^_p_=.016. There was no main effect of weight status, F(1, 49)=0.01, p=.911, η^2^_p_<.001. The valence x weight status interaction was not significant, F(1, 49)=0.40, p=.533, η^2^_p_=.008.

The P2 in response to the picture cues was analysed between 200 and 250ms after cue onset (see Figure 5). There was a main effect of valence, F(1, 49)=6.30, p=.015, η^2^_p_=.114. Pleasant picture cues (M=0.31 μV, SD=1.13) elicited a larger P2 than neutral picture cues (M=0.02 μV, SD=1.39). There was no significant main effect of weight status, F(1, 49)=0.02, p=.879, η^2^_p_<.001. There was a significant three-way valence x weight status x hemisphere interaction, F(2, 98)=3.90, p=.023, η^2^_p_=.074, but in the interaction between valence and weight status was not significant in either hemisphere (left hemisphere: F(1, 49)=3.55, p=.065, η^2^_p_=.068; midline: F(1, 49)=0.62, p=.434, η^2^_p_=.013; right hemisphere: F(1, 49)=0.26, p=.611, η^2^_p_=.005).

##### 3.3.1.3 N2 – no significant effects

For the taste cues, the N2 was analysed between 250 and 320ms after cue onset over electrodes Fp1/2, AF7/8, F7/8, FT7/8 (see Figure 6). We found no main effect of valence, F(1, 49)=2.22, p=.142, η^2^_p_=.043, weight status, F(1, 49)=0.47, p=.498, η^2^_p_=.009, nor their interaction, F(1, 49)=0.03, p=.876, η^2^_p_=.001.

For the picture cues, the N2 was also analysed between 250 and 320ms after cue onset (see Figure 6). We found no main effect of valence, F(1, 49)=1.34, p=.253, η^2^_p_=.027, weight status, F(1, 49)=2.40, p=.128, η^2^_p_=.047, nor theirinteraction F(1, 49)=0.04, p=.847, η^2^_p_=.001.

##### 3.3.1.4 P3 – effects of taste valence

For the taste cues, the P3 was analysed between 420 and 550ms after cue onset at electrodes P3/z/4 and PO7/z/8 (see Figure 7). We found a significant main effect of valence, F(1, 49)=4.21, p=.046, η^2^_p_=.079. P3 mean amplitude was more positive to pleasant (M=2.57 μV, SD=2.36) compared to neutral taste cues (M=2.14 μV, SD=2.07). There was no main effect of weight status, F(1, 49)=2.47, p=.122, η^2^_p_=.048, nor interaction between valence and weight status, F(1, 49)=1.24, p=.271, η^2^_p_=.025.

For the picture cues, the P3 was analysed between 350 and 420ms after cue onset at the same electrodes (see Figure 7). We found no main effect of valence, F(1, 49)=1.42, p=.238, η^2^_p_=.028, weight status, F(1, 49)=0.02, p=.903, η^2^_p_<.001, nor their interaction F(1, 49)=0.42, p=.520, η^2^_p_=.009.

##### 3.3.1.5 Stimulus-preceding negativity (SPN) – effects of taste valence

For the taste cues, the SPN (see Figure 6) was analysed at electrodes Fp1/2, AF7/8, F7/8, and FT7/8, between 480 and 750ms after cue onset (the 270ms preceding delivery of the taste stimulus). We found a significant main effect of valence, F(1, 49)=9.11, p=.004, η^2^_p_=.157. SPN was more negative for pleasant (M=-1.64 μV, SD=2.75) than neutral taste cues (M=-0.87 μV, SD=1.81). There was no main effect of weight status, F(1, 49)=0.24, p=.630, η^2^_p_=.005nor significant interaction between valence and weight status, F(1, 49)=0.72, p=.401, η^2^_p_=.014.

The SPN in response to the picture cues (Figure 6) was also analysed between 480 and 750ms (the 270ms preceding presentation of the pictures) at electrodes Fp1/2, AF7/8, F7/8, and FT7/8. There was no significant main effect of valence, although the p-value does indicate a trend, F(1, 49)=4.01, p=.051 η^2^_p_=.076. There was no main effect of weight status, F(1, 49)<0.01, p=.974, η^2^_p_<.001, nor significant interaction between valence and weight status, F(1, 49)=0.30, p=.584, η^2^_p_=.006.

#### 3.3.2 Target-elicited ERPs

##### 3.3.2.1 P1 – no significant effects

The P1 in response to the taste stimuli was analysed at electrodes TP7/8, P9/10, PO7/8, and O1/2, between 100 and 180ms (see Figure 8). There was no main effect of valence, F(1, 49)=0.95, p=.335, η^2^_p_=.019, or weight status, F(1, 49)<0.01, p=.998, η^2^_p_<.001nor their interaction, F(1, 49)=3.12, p=.083, η^2^_p_=.083.

The P1 in response to the picture stimuli was analysed at electrodes TP7/8, P9/10, PO7/8, and O1/2, between 150 and 190ms (see Figure 8). There was no main effect of valence, F(1, 49)=0.01, p=.938, η^2^_p_<.001, weight status, F(1, 49)=0.02, p=.894, η^2^_p_=<.001nor their interaction, F(1, 49)=2.01, p=.163, η^2^_p_=.039.

##### 3.3.2.2 N1 – no significant effects

The N1 in response to the taste stimuli was analysed between 180 and 250ms at electrodes TP7/8, P9/10, PO7/8, and O1/2 (see Figure 8). There was no main effect of valence, F(1, 49)=1.00, p=.322, η^2^_p_=.020 or weight status, F(1, 49)=0.11, p=.744, η^2^_p_=.002nor their interaction, F(1, 49)=3.65, p=.062, η^2^_p_=.069.

In response to the picture stimuli, the N1 was analysed at electrodes TP7/8, P9/10, PO7/8, and O1/2, between 190 and 240ms (see Figure 8). There was no main effect of valence, F(1, 49)=3.39, p=.072, η^2^_p_=.065, weight status, F(1, 49)=0.61, p=.440, η^2^_p_=.012, nor their interaction, F(1, 49)=0.07, p=.796, η^2^_p_=.001.

##### 3.3.2.3 P2 – no significant effects

The P2 in response to the taste stimuli was analysed at electrodes P9/10, PO7/8, and O1/2 between 260 and 320ms (see Figure 9). There was no main effect of valence, F(1, 49)<0.01, p=.958, η^2^_p_<.001, weight status, F(1, 49)<0.01, p=.986, η^2^_p_<.001, nor their interaction, F(1, 49)=3.91, p=.054, η^2^_p_=.074.

For the picture stimuli, the P2 was analysed at electrodes P9/10, PO7/8, and O1/2 between 260 and 350ms (see Figure 9). There was no main effect of valence, F(1, 49)=0.39, p=.535, η^2^_p_=.008, weight status, F(1, 49)=0.24, p=.624, η^2^_p_=.005, nor their interaction, F(1, 49)=0.04, p=.841, η^2^_p_=.00

##### 3.3.2.4 N2 – no significant effects

We analysed the N2 in response to the taste stimuli at electrodes AF3/z/4, F1/z/2, and FC1/z/2, between 250 and 320ms (see Figure 10). Their was no main effect of valence, F(1, 49)=2.07, p=.156, η^2^_p_=.041, weight status, F(1, 49)=0.04, p=.846, η^2^_p_=.001, nor their interaction, F(1, 49)=1.81, p=.185, η^2^_p_=.036.

For the picture stimuli, the N2 was analysed between 250 and 350ms at electrodes AF3/z/4, F1/z/2, and FC1/z/2 (see Figure 10). There was no main effect of valence, F(1, 49)<0.01, p=.950, η^2^_p_<.001, weight status, F(1, 49)=0.70, p=.408, η^2^_p_=.014, nor their interaction, F(1, 49)=1.33, p=.254,η^2^_p_=.027.

##### 3.3.2.5 P3 – no significant effects

We analysed the P3 in response to the taste stimuli at electrodes TP7/8, P9/10, PO7/8, and O1/2, between 420 and 500ms (see Figure 8). There was no main effect of valence, F(1, 49)=1.77, p=.190, η^2^_p_=.035, weight status, F(1, 49)=0.03, p=.874, η^2^_p_=.001, nor their interaction, F(1, 49)=0.56, p=.458, η^2^_p_=.011.

For the picture stimuli, the P3 was analysed at electrodes TP7/8, P9/10, PO7/8, and O1/2 between 400 and 480ms (see Figure 8). There was no main effect of valence, F(1, 49)=0.14, p=.708, η^2^_p_=.003, weight status, F(1, 49)=0.22, p=.640, η^2^_p_=.004, nor their interaction, F(1, 49)=0.01, p=.930, η^2^_p_<.001.

### 3.4 Exploratory whole-brain EEG analyses

In addition to the literature-driven analyses, we ran exploratory whole-brain analyses on both epochs to determine whether any effects were observed outside of the predicted effects. This was motivated by the relative lack of a priori studies testing for ERP differences as a function of weight status.

#### 3.4.1 Exploratory findings in the cue-elicited epoch

In response to the taste cues, we found one significant cluster showing a main effect of valence. This right-frontal cluster (k=1065, p=.035) was observed over the AF4 electrode and showed significant peaks between 520 and 575ms after cue onset (all Fs(1, 98)≥9.23, all ps≤.001). These peaks showed a more negative deflection in response to the pleasant taste cues compared to the neutral taste cues. No significant clusters were found for the interaction between weight status and valence.

In response to the picture cues, we also found one significant cluster showing a main effect of valence. This cluster (k=2844, p=.001) occurred centro-posteriorly, extending into the left and right hemispheres over the CP3, CP1, and Pz electrodes, and contained significant peaks between 680 and 690ms (both Fs(1, 98)≥18.33, both ps<.001). These peaks showed a more positive deflection in response to the pleasant picture cues compared to the neutral picture cues. No significant clusters were found for the interaction between weight status and valence.

#### 3.4.2 Exploratory findings in the target-elicited epoch

In response to the tastes, we found one significant cluster showing a main effect of valence. This cluster (k=1497, p=.032) occurred fronto-centrally, slightly right of the midline over electrodes FC2 and FC4. The cluster contained two significant peaks between 885 and 960ms (both Fs(1, 98)≥13.02, both ps<.001). The peaks showed a more positive deflection in response to the pleasant tastes compared to the neutral tastes. No significant clusters showing a weight status by valence interaction were found. Moreover, no significant clusters were found in response to the pictures.

## 4. Discussion

This study investigated whether normal-weight and overweight individuals differ in their processing of pleasant taste stimuli and cues indicating their imminent presentation. Pictures were also utilised to determine whether any weight-group differences were specific to food or were general differences in response to cues or stimuli with affective value. The manipulation of taste valence was successful, evident in both ERP and behavioural measures. Pleasant taste cues elicited larger P3 and SPN amplitude; the delivery of pleasant taste modulated EEG signal in a whole-brain exploratory analysis; and participants rated the sweet taste more favourably than the neutral taste. In agreement with our hypothesis, results showed a weight-group difference in pleasantness and comparative pleasantness specific to taste stimuli, such that overweight participants rated pleasant stimuli as less pleasant than normal-weight participants, and as less pleasant than expected. However, our hypothesis that the interaction between weight-group and valence would modulate ERP components was not supported. Rather, there were weight-related neural differences in early attention to cues.

### 4.1. Valence effects confirm conditioning procedure success

We observed a main effect of valence on a number of ERPs and behavioural measures, confirming that our conditioning procedure was successful. Pleasant taste and picture stimuli received significantly higher liking ratings than neutral stimuli, a result that remained significant in each group (see below for discussion of the interaction of valence and weight status). For cue-elicited ERPs, P3 amplitude was more positive in response to pleasant than neutral taste cues. The P3 is associated with consciously attending to salient or personally-relevant stimuli, and updating one’s mental representation of them (Polich, 2007; Ullsperger et al., 2014). This therefore suggests participants paid more sustained attention to pleasant taste cues, perhaps because they considered them more motivationally salient (Franken et al., 2011). This effect of valence on P3 amplitude was found for taste but not picture cues, suggesting that participants may not have found pleasant pictures more motivationally salient. Both the valence and arousal properties of visual stimuli have been found to independently affect the amplitude of early, mid-latency, and late ERPs (Olofsson et al., 2008; Recio, Conrad, Hansen, & Jacobs, 2014). Therefore, the null effect may be in part because although participants rated pleasant pictures as more liked than neutral ones, intensity ratings were higher for neutral than pleasant pictures. This difference occurred despite effort to match the pleasant and neutral pictures on their arousal level based on the IAPS arousal ratings.

Valence also modulated SPN amplitude and a corresponding cluster in the exploratory analysis, both of which were more negative in response to pleasant than neutral taste cues. The timing of the SPN is tied to presentation of the target stimulus, and is an index of attentional engagement and motor preparation during the anticipation of this target stimulus (Hirao, Murphy, & Masaki, 2016). Cues indicating immenent delivery of the pleasant taste thus engaged greater attention and expectation (Fuentemilla et al., 2013). This converges with the enhanced P3 amplitude by pleasant taste cues.

Responses to taste stimuli themselves did show a modulation by valence in the exploratory whole-brain analysis at a fronto-central midline cluster between 885 and 960ms. Mean voltage was more positive for pleasant than neutral taste stimuli, indicating increased sustained attention when tasting the pleasant stimuli. Franken et al. (2011), who used the same neutral control stimulus we used, found a similar positive potential occurring relatively late after taste delivery that was more positive in response to a sweet, pleasant taste compared to their control stimulus. However, valence did not significantly modulate any of the ERP components time-locked to taste delivery.

Picture cue valence also modulated ERPs. P2 amplitude was greater for pleasant than neutral picture cues. This suggests pleasant picture cues elicited more preconscious attention than neutral picture cues (Franken et al., 2011). P2 amplitude enhancement is observed for stimuli with higher motivational salience (Wang, Kleffner, Carolan, & Liotti, 2018). The exploratory analysis also revealed a cluster at centro-parietal midline electrodes, around 680ms after the presentation of the picture cues, which was more positive for pleasant than neutral picture cues. A similar late positive potential at this scalp location is often found in response to emotional stimuli, with more positive mean voltage in response to emotionally salient (i.e. pleasant or unpleasant) compared to neutral stimuli (Hajcak, Dunning, & Foti, 2009; Olofsson, Nordin, Sequeira, & Polich, 2008; Schupp et al., 2000). This late positive activity is an index of sustained allocation of attention, occurring in particular to motivationally-relevant stimuli (Cona, Kliegel, & Bisiacchi, 2015). Participants therefore showed stronger sustained attention to cues predicting the presentation of pleasant pictures. These findings suggest there were differences in allocation of attention to the cues depending on the valence of the picture that they were paired with.

Taken together, our ERP results suggest that cue valence modulates processing, but does so at different stages for taste and picture cues. The results validate our manipulation because they show that participants paid selective attention to cues depending on the valence of the target those cues were paired with.

### 4.2. Weight-group differences in ERP indices of cue processing

To test whether weight status was associated with changes in early perceptual and pre-conscious attentional processing of cues, we analysed cue-elicited N1 and P2 amplitude. N1 amplittude was larger in the normal-weight group in response to both taste and picture cues, regardless of cue valence. This suggests the normal-weight group showed greater early allocation of attention to cues than the overweight group, and that this difference was not food-specific (Flores et al., 2015; Kemmotsu & Murphy, 2006). The interpretation that reduced N1 amplitude indicated the overweight group paid less attention to the cues was supported by the behavioural data, which showed reduced learning from the cues in this group. There was no effect of weight group on P2 amplitude. To determine whether weight status is associated with changes in later, conscious attention or expectation we also examined N2, P3 and SPN amplitude. No differences between the two groups were found.

These results are consistent with models of obesity that suggest that overeating is associated with changes in attention and executive function (Cserjesi et al., 2009 (Sabia et al., 2009; Wolf et al., 2007). Some previous studies have shown that BMI is negatively correlated with executive function (Sabia et al., 2009; Wolf et al., 2007). Increased BMI has also been associated with greater distractability and reduced capacity for sustained attention (Cserjesi et al., 2009).

A caveat for this interpretation is that the normal-weight group was significantly younger than the overweight group. One study investigating age-related decline in visual attention found older adults showed reduced anterior N1 amplitude in response to visual stimuli (Wiegand et al., 2014). However, other studies looking at age-related changes to the N1 have either not found any effects of age (Polich, 1997), or found N1 amplitude increasing with age during adulthood (De Blasio & Barry, 2018; Reuter et al., 2019). Our results remained significant when we partialled out the effects of age, suggesting that the age difference between our two groups did not cause the group differences in cue-elicited N1 amplitude.

### 4.3 Interaction between valence and weight status

Whilst both overweight and normal-weight participants found the pleasant taste stimulus more pleasant than neutral, this difference was significantly smaller for the overweight group. Despite their increased level of hunger, the overweight group rated the pleasantness of pleasant and neutral taste stimuli as more similar than did the normal-weight group. This could indicate the overweight group were less sensitive to the difference between the two taste stimuli pointing to blunted sensory processing of the gustatory stimuli. Alternatively, the overweight group may differentiate less between the hedonic values of the two stimuli, with sensory processing intact. The lack of significant group differences in intensity ratings of the taste stimuli suggests the latter is more likely. Rather, the data are consistent with the proposal that overweight individuals exhibit a change in hedonic perception of taste (Devoto et al., 2018; Stice & Burger, 2019).

This difference in hedonic rating of the taste stimuli may be driven by weight-group differences in cue processing. The comparative pleasantness ratings are one way to test this. The neutral taste stimulus was always the same, thus as trials progressed, participants could learned exactly what to expect. This was the case for normal-weight participants, where comparative pleasantness ratings approximated “as pleasant as expected” as trials progressed. This learning did not occur in overweight participants.

We predicted, based on the previous literature, that the electrophysiological data ouldw show enhanced food reward anticipation in the overweight group (Devoto et al., 2018; García-García et al., 2013; Hume et al., 2015; Lawrence et al., 2012; Nijs et al., 2010; Stice et al., 2008; Stice & Burger, 2019). However, contradicting this hypothesis, the interaction between valence and weight group was not significant in any of the ERP components, neither in response to cues or to outcome delivery, nor in the whole-brain exploratory analysis.

These results do not support the proposal that overweight people experience a heightened anticipation in response to food cues (Hume et al., 2015; Nijs et al., 2008; Stice & Burger, 2019; Volkow et al., 2013). Our data also do not support models of obesity that argue overeating is related to a reduction in experienced reward from food (Volkow et al., 2013a; Volkow et al., 2013b; Wang et al., 2001) nor those proposing reduced sensory processing of food (Bartoshuk, Duffy, Hayes, Moskowitz & Snyder, 2006).

### 4.4. Limitations

We chose to study neural responses during satiety, as prior research has suggested this is the state in which weight-driven differences are strongest (Nijs et al., 2010). Therefore, we instructed our participants to eat before the experiment, to ensure they ate as much as they usually would for satiation, rather than reducing intake due to social desireability in the lab, and to avoid differences in satiation induced by a set meal size. We took every effort to ensure satiation, with repeated instructions and time given to eat. Nevertheless, the overweight group were more hungry than the normal-weight group when they entered the lab, and craved more food at the end of the conditioning task. Notably, there were no differences between the groups immediately before the conditioning task and no differences in hunger at the end of the conditioning task. While the group differences in food craving may have influenced our results, it is unlikely that group differences in hunger would eliminate the influence of weight on the electrophysiological data.

Our lack of differences between the overweight group and the normal-weight group goes against a number of fMRI studies that reported reduced activation in overweight people in response to pleasant taste stimuli (Babbs et al., 2013; Green et al., 2011; Stice & Burger, 2019; Stice et al., 2008). One methodological factor that might explain these divergent findings is the type of pleasant taste stimulus used. This study used two types of squash (i.e. a sugary liquid) as the pleasant taste. In other studies, a common pleasant stimulus was a milkshake, which contains fat as well as sugar. It might be that these differences between the weight groups only present themselves in response to fatty foods specifically, or to foods containing a combination of fat and sugar, and not in response to sugar alone. Indeed, the findings of a recent fMRI study suggest that consuming food that is both high in fat and high in carbohydrates induces greater dorsal striatal activity than high-fat or high-carbohydrate food alone (DiFeliceantonio et al., 2018). In this study, participants were also more willing to pay to receive food that was both high in fat and high in carbohydrates compared food that was only high-fat or high-carbohydrate. These findings are in line with previous findings from animal studies suggesting that the combination of fat and carbohydrates in food induces more excessive eating in rats than food that is high in only one of these macronutrients (Lucas & Sclafani, 1990). To the author’s knowledge, no studies thus far have compared overweight and normal-weight people’s neural responses to foods containing different macronutrients. This could therefore be a focus point for future studies.

A caveat of our interpretations thus far is that the pleasantness ratings for the neutral taste stimuli, on a scale from one to nine, were not rated as approximately five (the midpoint of the scale), which would be entirely neutral. Instead, on average the neutral stimulus was rated as slightly aversive, potentially changing the interpretation of the results of our study so that the comparison here is not between pleasant and neutral taste stimuli, but to pleasant and aversive stimuli. A follow-up study confirmed that this slight aversiveness was not due to a misunderstanding of the nine-point pleasantness scale (Baines et al., 2020). Rather, participants rated the nominally “neutral” stimulus used here as mildly aversive. Interpretation of the ERP results should therefore consider a comparison between pleasant and mildy aversive, rather then neutral, stimuli.

The purpose of this study was to investigate whether any hypothesised differences between overweight and normal-weight participants in response to food and food cues were due to early perceptual differences in processing, or differences in late, cognitive mental processes. The key difference in overweight and normal-weight participants in our data was a reduction in early allocation of attention to cues. This was not specific to food, but also evident in response to picture cues. Comparative pleasantness ratings also suggested an inability to learn from the taste cues. Our data therefore suggest a difference in cue-related learning and attention in overweight. This may be an important factor in disordered eating, when consumption becomes dissociated with physiological hunger signals.

## Footnotes

1 The following IAPS pictures were used in the pleasant picture condition: 1340, 1440, 1460, 1463, 1540, 1710, 1722, 1750, 1811, 1920, 1999, 2040, 2050, 2058, 2070, 2071, 2080, 2091, 2209, 2260, 2300, 2314, 2352, 2550, 2630, 4532, 4599, 4603, 4610, 4612, 4614, 4624, 4628, 4641, 5199, 5210, 5260, 5270, 5460, 5470, 5480, 5600, 5660, 5700, 5814, 5825, 5829, 5830, 5833, 5910, 5994, 7502, 7508, 8162, 8170, 8420, 8499, 8500, 8503, 8531.

2 The following IAPS pictures were used in the neutral picture condition: 1121, 1122, 1302, 1303, 1310, 1313, 1350, 1390, 1560, 1645, 1726, 1820, 1908, 1945, 2018, 2122, 2210, 2211, 2220, 2372, 2458, 2616, 2661, 2681, 2704, 2780, 2810, 3005.2, 3310, 5455, 5920, 5950, 6837, 6900, 6910, 6930, 7077, 7211, 7496, 7497, 7504, 7505, 7560, 7600, 7620, 7632, 7640, 8211, 8250, 8251, 8260, 8341, 8475, 8620, 9080, 9402, 9411,9422, 9468, 9582.

